# Mesoscale chromatin confinement facilitates target search of pioneer transcription factors in live cells

**DOI:** 10.1101/2024.07.18.604200

**Authors:** Zuhui Wang, Bo Wang, Di Niu, Chao Yin, Ying Bi, Claudia Cattoglio, Kyle M Loh, Luke D Lavis, Hao Ge, Wulan Deng

## Abstract

Pioneer transcription factors (PTFs) possess the unique capability to access closed chromatin regions and initiate cell fate changes, yet the underlying mechanisms remain elusive. Here, we characterized the single-molecule dynamics of PTFs targeting chromatin in living cells, revealing a notable “confined target search” mechanism. PTFs like FOXA1, FOXA2, SOX2, OCT4 and KLF4 sampled chromatin more frequently than non-pioneer factor MYC, alternating between fast free diffusion in the nucleus and slower confined diffusion within mesoscale zones. Super-resolved microscopy showed closed chromatin organized as mesoscale nucleosome-dense domains, confining FOXA2 diffusion locally and enriching its binding. We pinpointed specific histone-interacting disordered regions, distinct from DNA-binding domain, crucial for confined target search kinetics and pioneer activity within closed chromatin. Fusion to other factors enhanced pioneer activity. Kinetic simulations suggested transient confinement could increase target association rate by shortening search time and binding repeatedly. Our findings illuminate how PTFs recognize and exploit closed chromatin organization to access targets, revealing a pivotal aspect of gene regulation.

## Introduction

Cell fate transitions rely on specific transcription factors (TFs) to activate new gene expression programs. However, the presence of nucleosomes and compaction of closed chromatin hinder most TFs and the transcription machinery from accessing DNA templates^1,2^. Pioneer transcription factors (PTFs), such as FOXA factors, exhibit remarkable abilities to access DNA targets even when nucleosome-bound, facilitating the opening of closed chromatin and triggering cell fate changes^3–6^. Despite extensive studies focusing on the structural and functional aspects of PTFs binding to nucleosomes^6–12^, the precise mechanisms underlying their initial search for and binding to closed chromatin, setting PTFs apart from other TFs, remains unclear.

Single-molecule localization microscopy enables the direct visualization of individual molecules in cells with spatial resolution surpassing the diffraction limit^13^. Live-cell single-molecule tracking (SMT) has revealed dynamic interactions between TFs^14–16^ or transcription machinery^17,18^ with chromatin, challenging the notion of stable protein complexes on chromatin^19,20^. Understanding TF kinetics in the genome-wide target search is crucial for deciphering transcription regulation^21,22^. Eukaryotic TFs primarily utilize 3D diffusion in the nucleus, but it is unclear if they employ additional mechanisms for efficient chromatin target search, especially in closed chromatin where PTFs operate. Prokaryotic TFs employ local sampling of DNA, known as facilitated diffusion, alongside the 3D diffusion to enhance target search efficiency^23,14^. While facilitated diffusion may occur in the open chromatin of eukaryotic cells^24,16^, it falls short in explaining target search in closed chromatin, densely packed with nucleosomes. Despite studies on PTF kinetics^25–28^, questions remain about mechanism of efficient target binding in closed chromatin.

In this study, we combined live-cell fast SMT, fixed-cell multi-color PALM, computational simulation, kinetic modeling, and epigenomic profiling to elucidate the dynamic process of PTFs searching for targets on chromatin in living cells. We revealed a notable “confined target search” mechanism that enables PTFs to access targets in closed chromatin. In this mechanism, the PTF alternates between fast free diffusion in the nucleus and slower confined diffusion within mesoscale compacted chromatin domains in closed chromatin. This behavior, facilitated by the interaction of their disordered regions with nucleosomes, enables shorter search time and repetitive binding trials to nucleosomal targets. Our findings suggest that the physical organization of chromatin influences TF kinetics in living cells and that TFs exploit local chromatin organization to enhance their activities.

## Main text

### Fast spaSMT reveals frequent chromatin binding by PTFs

To capture both bound molecules with low mobility and those in fast diffusion in living cells, we set up fast stroboscopic photo-activation SMT (fast spaSMT) microscopy at millisecond (ms) temporal resolution (Fig. 1, A and B). This technique uses highly inclined and laminated optical (HILO) sheet illumination ^29^ to enhance the signal-to-noise ratio of imaging individual molecules in the nucleus. The protein of interest is tagged with HaloTag (Halo for short) and labelled with photoactivatable Janelia Fluor dyes^30^ (Supplementary Fig. 1A). In living cells, single fluorophores are then sparsely and stochastically activated by a 405 laser and recorded with stroboscopic illumination to obtain sharp images. A large population of single-molecule trajectories are generated and analyzed with the kinetic modeling algorithm Spot-On ^31^ to infer chromatin-bound fractions and diffusion constants.

**Figure 1.**
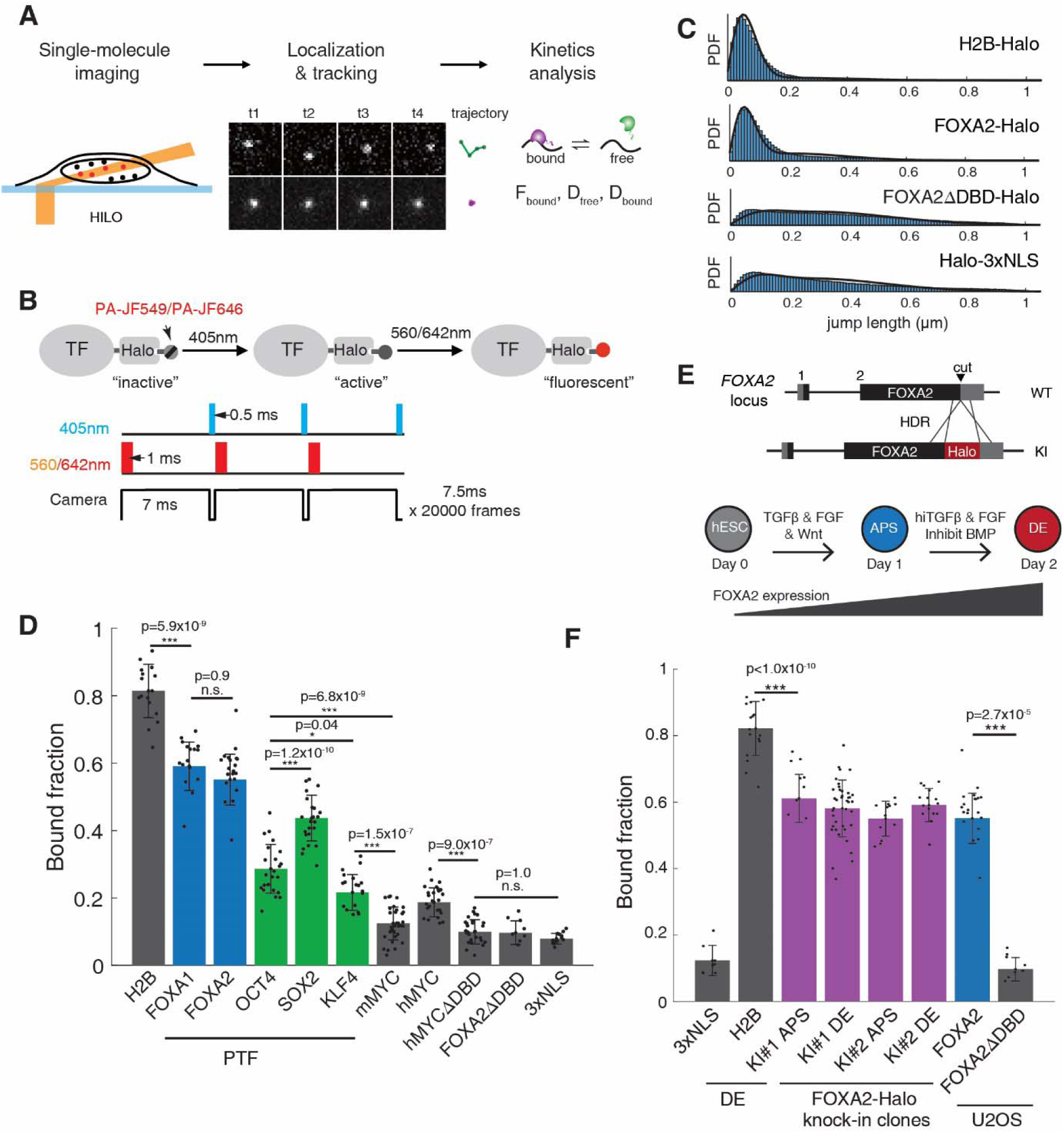
PTFs sample chromatin frequently for targets by fast spaSMT. (**A**) The schematic diagram of single-molecule tracking pipeline. (**B**) The schematic diagram of fast spaSMT with stroboscopic illumination and photoactivable fluorophore PA-JF549 or PA-JF646. (**C**) The probability distribution function of jump length from all pooled trajectories of indicated factors at 133 Hz imaging frequency. (**D**) Bound fractions (mean ± S.D., dots represent data points) derived from two-state Spot-On analyses on fast spaSMT trajectories of indicated Halo-tagged factors exogenously expressed in MEF cells (murine OCT4, SOX2, KLF4, and MYC) or U2OS cells (human FOXA1, FOXA2, MYC, DBD deletion mutants, H2B and 3xNLS) with Tet-on system. Colored molecules are PTFs. (**E**) (Top) Illustration of CRISPR/Cas9 mediated knock-in (KI) of HaloTag to the C-terminus of the endogenous FOXA2 locus in hESCs. (Bottom) Differentiation of hESC to anterior primitive streak (APS) and definitive endoderm (DE). FOXA2 is not expressed in hESC and induced to express upon differentiation. (**F**) Bound fractions (mean ± S.D., dots represent data points) of indicated factors in following cell conditions: stable hESC cell lines expressing Halo-tagged 3xNLS or H2B were induced to DE state; two FOXA2-Halo homozygous knock-in clones were induced to APS or DE state; U2OS cells expressed FOXA2 variants. Significance was calculated using one-way ANOVA followed by Tukey’s multi-comparison, *p-value<0.05, **p-value<0.01, ***p-value<0.001, n.s., not significant. (**C, D, F**) n=16-58 biologically independent cells for each sample, see Supplementary Table 1.

We first used fast spaSMT to investigate a number of pioneer factors in comparison to non-pioneer factors. The examined PTFs included FOXA1^3^, FOXA2^32^, and reprogramming factors OCT4, SOX2, and KLF4, while MYC was recognized as a non-pioneer factor^4^. Murine OCT4, SOX2, KLF4 and MYC were expressed in mouse embryonic fibroblasts (MEFs), and human FOXA1, FOXA2 and MYC were expressed in human bone osteosarcoma U2OS cells. These cells lacked endogenous expression and the Halo-tagged fusion proteins were exogenously expressed to a moderate level with the Tet-on inducible system. Additionally, freely diffusing nuclear localization signal (NLS) and core histone H2B were assessed in U2OS cells as controls. A substantial number of fast spaSMT trajectories (10^5^ to 10^6^) were collected per factor, and subsequently analyzed by Spot-On with two-state kinetic modeling (Fig. 1, C and D; Supplementary Fig. 1B; Supplementary Video 1; Supplementary Table 1) or vbSPT^33^ (Supplementary Table 2). As anticipated, the majority of H2B (81.4%) and a minor fraction of NLS (7.9%) were found to be bound to chromatin. While the non-pioneer factors mMYC and hMYC showed a low chromatin-bound fraction at 12.5% and 18.7% respectively, the PTFs exhibited higher values, with FOXA factors the highest (FOXA1, 59.1% and FOXA2 55.1%) even with varying protein expression levels (Supplementary Fig. 1C). Deletion of the DNA-binding domain (DBD) reduced the bound fraction to background levels, indicating that these PTFs frequently bound chromatin to sample for cognate targets.

FOXA2 plays pioneering roles in formation of endoderm lineages^32,34,35^. To further elucidate the dynamics of endogenous PTF-chromatin interactions, we generated homozygous FOXA2-Halo knock-in (KI) human embryonic stem cell (hESC) clones and differentiated them to anterior primitive streak (APS) and subsequently definitive endoderm (DE)^36^ (Fig. 1E). Throughout this process, FOXA2 was rapidly induced. Importantly, the introduction of the Halo did not interfere with FOXA2 expression or DE differentiation (Supplementary Fig. 1, D to H; Supplementary Video 2). We next induced the KI cells to APS and DE stages, capturing 1.8×10^5^ and 3.9×10^5^ fast spaSMT trajectories of FOXA2-Halo, respectively. The results consistently revealed a substantial fraction of FOXA2 molecules (55-61%) in the chromatin-bound state in both APS and DE cells, as observed in two independent FOXA2-Halo KI clones (Fig. 1F; Supplementary Fig. 1I), aligning with the findings from U2OS cells. Therefore, these results indicated that the PTF molecules frequently interacted with chromatin.

### PTFs exhibit locally confined diffusion on chromatin

To further investigate the chromatin sampling mechanism of PTFs, we analyzed the directional preference of molecule diffusion^37,38^ (Fig. 2A). Specifically, we used a hidden Markov model (HMM) to classify segments (displacement between two sequential localizations) of molecule trajectories into bound and free states and measured the angles between two consecutive free segments. We found that all examined PTFs exhibited markedly anisotropic diffusion, with a higher probability to move backward than other directions (peaking around 180°) (Supplementary Fig. 2, A and B). We quantified the degree of anisotropic diffusion by the “fold anisotropy”, *f*_180/0_, and found that diffusion of PTFs showed generally high anisotropy (Fig. 2B). In contrast, nuclear proteins including non-pioneer factors MYC showed typical isotropic diffusion. Deletion of FOXA2 DBD reduced the degree of anisotropic diffusion to background levels, indicating that the observed anisotropic diffusion was associated with DNA binding activities.

**Figure 2.**
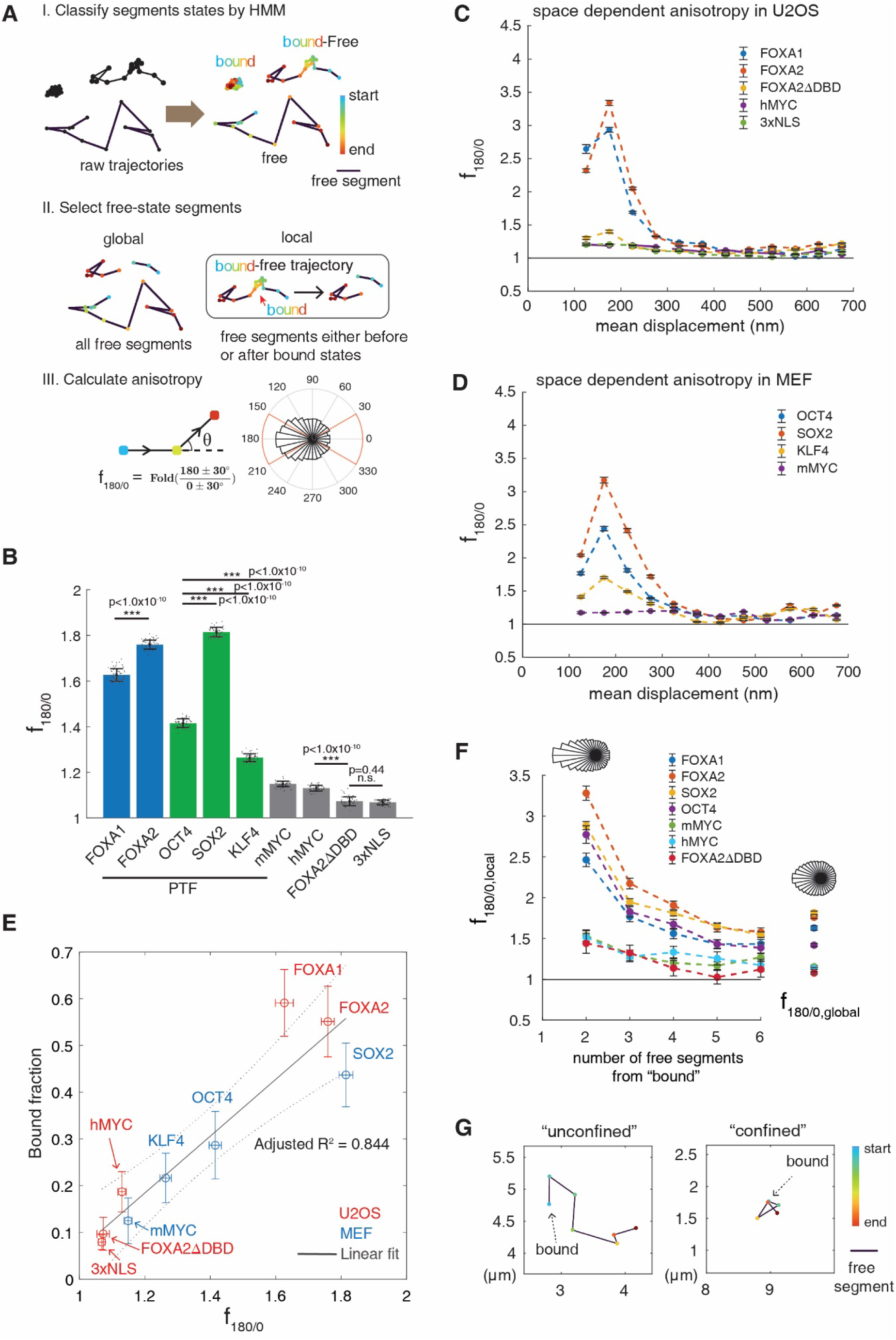
PTFs exhibit anisotropic diffusion and confined target search on chromatin. (**A**) Illustration of the analysis pipeline of global and local diffusion anisotropy. The molecule localizations of exemplary trajectories are color-coded illustrating start to end, and the free segments are particularly colored black. (**B**) *f_180/0_* (mean ± S.D., dots represent data points) of indicated molecules calculated from all free segments (global anisotropy) in U2OS cells and MEF cells. Colored molecules are PTFs. (**C** and **D**) Space-dependent anisotropy examined by *f_180/0_* against mean displacement length of selected free segments. Error bars show standard errors from 50 subsamplings of segment angles with 50% replacement. (**E**) Plots of global anisotropy *f_180/0_*versus bound fraction of indicated molecules. Horizontal and vertical error bars are standard deviations of *f_180/0_* and bound fraction, with the center denoting the mean. Gray solid line represents the linear regression, and gray dotted line represents the 95% confidence interval of the regression. The adjusted R-square is calculated by adjusting the number of data points. (**F**) *f_180/0, local_* (mean ± S.D.) were calculated from a variable number of free segments before and after a binding event (local anisotropy, left) and compared to *f_180/0_* of all free segments (global anisotropy, right). The representative angle distribution histograms of FOXA2 are shown on the top. (**G**) Representative single-molecule trajectories showing diffusion in either a “confined” or “unconfined” manner after a binding event. Localizations and segments are colored as in (A). Significance was calculated using one-way ANOVA followed by Tukey’s multi-comparison, *p-value<0.05, **p-value<0.01, ***p-value<0.001, n.s., not significant. (**B-F**) n=13-58 biologically independent cells for each sample, see Supplementary Table 1.

We next examined the degree of anisotropic diffusion as a function of the length of displacements flanking the angle (Fig. 2, C and D; Supplementary Fig. 2, C and D). All PTFs showed clear displacement-dependent anisotropy, peaking at displacements around 200 nm. This indicated that PTFs diffused anisotropically only at short displacements but diffused isotropically at longer displacements, suggestive of transiently confined diffusion within zones around the 200 nm mesoscale. Deletion of DBD significantly reduced diffusion anisotropy at this scale (Fig. 2C). In contrast, diffusion of NLS and non-pioneer MYC showed little anisotropy at any length of displacements. These results suggested that the PTF molecules uniquely adopt two alternating modes of target search in cells: free diffusion in the nucleus (isotropic) and confined diffusion within mesoscale zones (anisotropic).

We found that the level of confinement positively correlated with the fraction of chromatin-bound molecules (Fig. 2E). The correlation of the two measurements suggested that confined diffusion actively facilitates frequent chromatin binding. To test this, we selected state-switching trajectories (containing both free and bound states in one trajectory) and calculated the anisotropy of consecutive free segments before or after a bound state in trajectories, defined as “local anisotropy” as opposed to the “global anisotropy” measured from all free segments (Fig. 2A). Interestingly, PTFs showed high level of local anisotropy, i.e., confined diffusion, surrounding their chromatin binding events (Fig. 2F; Supplementary Fig. 2E). By visualizing randomly selected state-switching trajectories, we observed that PTF molecules tended to diffuse around their previous bound sites (“confined” diffusion), while non-pioneer MYC molecules tended to travel away from their previous bound sites randomly (“unconfined” diffusion) (Fig. 2G; Supplementary Fig. 2, F and G; Supplementary Video 3). We also noticed that PTFs became un-confined after a few frames (7.5 ms per frame), indicating that these PTFs molecules were only transiently confined in the local chromatin zone rather than permanently trapped (Fig. 2F). These results suggested that confined diffusion of PTFs was closely associated with frequent chromatin binding.

### FOXA2’s confined diffusion mapped to histone-interacting CTD

We next investigated the cause and consequence of chromatin confinement by focusing on FOXA2. FOXA2 is composed of a well-structured Forkhead DNA binding domain (FKHD) and two intrinsically disordered regions (IDRs) known as the N-terminal domain (NTD) and the C-terminal domain (CTD) (Fig. 3A). Previous studies have demonstrated that the CTDs of murine FOXA1/2 directly interact with purified core histones and are required for nucleosome binding ^3^. Impairment of this interaction significantly affects mouse embryonic development, altering gene activation and chromatin opening^32^. However, it is unclear how mechanistically CTD-histone interaction promotes pioneer activity. We confirmed CTD-histone interactions by co-IP experiments using nuclear lysates. FOXA2 interacted with core histone H3 and H4, and deletion of its CTD abolished this interaction (Fig. 3B). Additionally, CTD alone or deletion of DBD exhibited weak interaction with core histones compared to full-length FOXA2, indicating that DNA binding enhances CTD-histone interactions.

**Figure 3.**
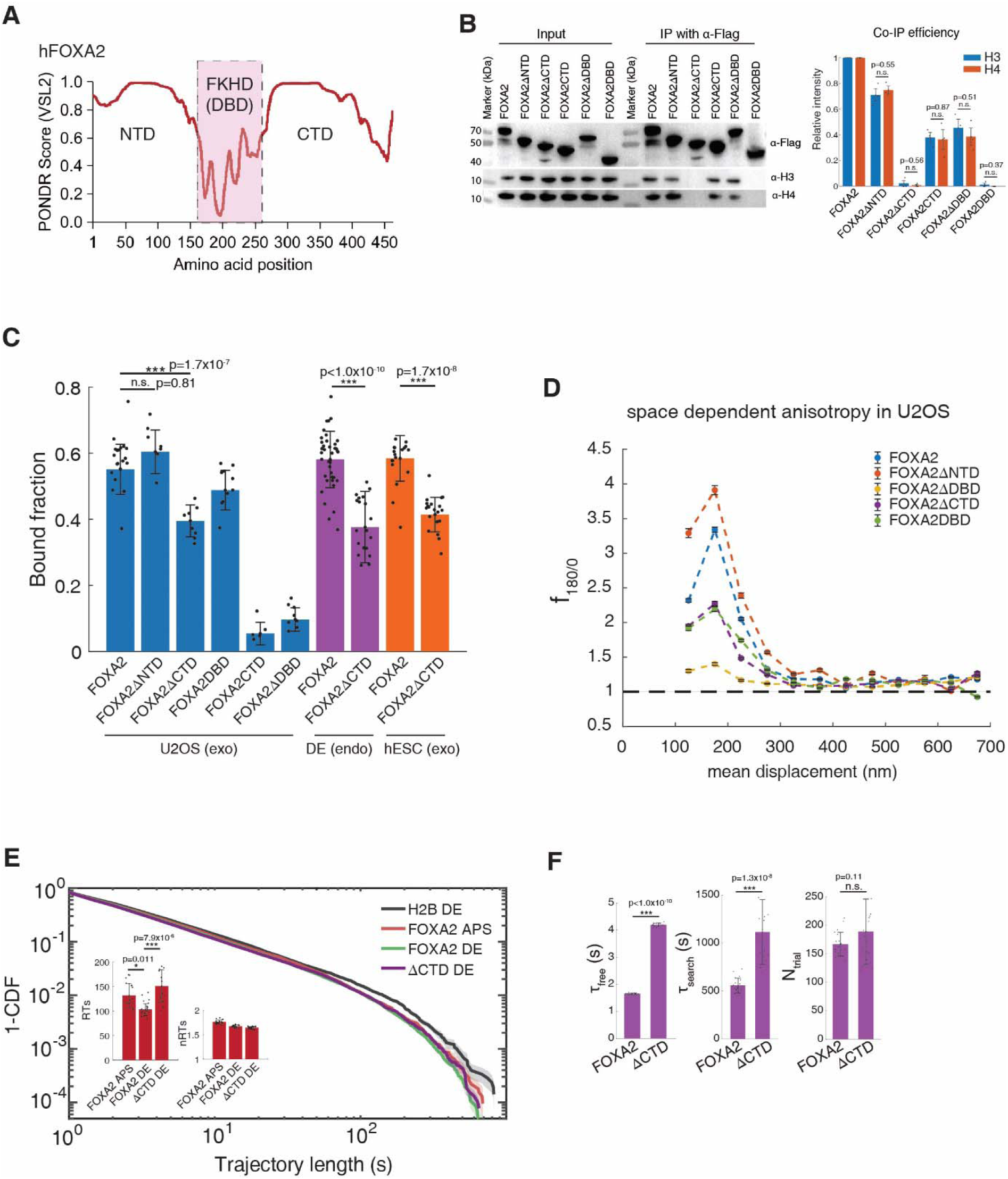
CTD_FOXA2_ is essential for FOXA2 anisotropic diffusion and target search. Predictor of naturally disordered regions (PONDR) scores of the human FOXA2 protein. (**B**) Co-IP using nuclear lysates of U2OS cells expressing Flag-tagged FOXA2 variants. Representative Western blots (left) and quantification (mean ± SEM, dots represent data points) of three replicates (right) are shown. n=3 biologically independent samples. (**C**) Bound fractions (mean ± S.D., dots represent data points) derived from two-state Spot-On analyses of fast spaSMT trajectories for indicated molecules in indicated cells. “exo” indicates exogenous expression, and “endo” indicates endogenous expression with knock-in clones. (**D**) Space-dependent anisotropy examined by *f_180/0_* against mean displacement length of selected free segments. Error bars show standard errors from 50 subsamplings of segment angles with 50% replacement. (**E**) Log-log survival curves of slowSMT trajectories of the indicated molecules (left). Insert: corrected specific and nonspecific residence times (RTs and RTns, mean±SEM, dots represent data points) of endogenously tagged FOXA2 or ΔCTD in DE cells. (**F**) Comparison of FOXA2 and ΔCTD target search kinetics. Results are mean ± S.D., dots represent data points. Significance was calculated using one-way ANOVA followed by Tukey’s multi-comparison in (**C** and **E**), two-sided t-test in (**B** and **F**), *p-value<0.05, **p-value<0.01, ***p-value<0.001, n.s., not significant. (**C-F**) n=16-58 biologically independent cells for each sample, see Supplementary Table 1.

We investigated the contribution of FOXA2 domains to its chromatin association dynamics using fast spaSMT. Deletion of DBD eliminated specific chromatin association, as expected for a TF (Fig. 3C). Remarkably, deletion of CTD significantly decreased the bound fraction of FOXA2 on chromatin (from 55.1% to 39.5%) (Fig. 3C) and also reduced confined diffusion (*f*_180/0, 200nm_ reduced from 3.34 to 2.27) in U2OS cells (Fig. 3D). The global and local anisotropy of its diffusion were similarly reduced (Supplementary Fig. 3, A to C). In comparison, deletion of NTD did not decrease but even slightly increased the chromatin bound fraction and diffusion anisotropy, suggesting inhibitory roles of NTD in chromatin association. Post-translation modification such as acetylation has been reported to attenuate FoxA1 nucleosome binding and remodeling binding^39^. Our focus remained on CTD. Therefore, we generated homozygous hESC clones with CTD deletion (ΔCTD-Halo) based on the FOXA2-Halo KI hESC clone (Supplementary Fig. 3D). Expression level of ΔCTD-Halo was the same as full-length FOXA2-Halo in stage-matched DE cells. Consistent with the results from U2OS cells, CTD deletion significantly reduced the chromatin-bound fraction (from 58.1% to 37.6%) (Fig. 3C) and confined diffusion in DE cells (Supplementary Fig. 3E).

Experiments with exogenously expressed FOXA2 and ΔCTD in hESC cells also produced consistent results (Fig. 3C and Supplementary Fig. 3F). These results indicated that CTD played essential roles in frequent chromatin association and confined diffusion of FOXA2.

We further explored the contribution of CTD to stable target binding using slowSMT. The slowSMT imaging module, with a long exposure time (500 ms) and low laser power, selectively captures stable-bound molecules while blurring fast-diffusing ones (Supplementary Video 4 and 5). By two-exponential fitting and photobleaching correction with H2B (Supplementary Fig. 3G), we derived the residence time of FOXA2 specific binding (RTs) around 2 minutes and that of non-specific binding (RTns) around 2 seconds (Fig. 3E). Notably, our results indicated that FOXA2 has an extended target residence time (RTs) than other reported TFs^20^, similar to that of GAGA factor ^28^. Importantly, deletion of CTD did not reduce RTs and RTns, showing that while CTD played essential roles in target searching, it played little roles in the stability of target binding. We observed a slight increase in RTs upon CTD deletion, which could be attributed to the ΔCTD favoring open chromatin and exhibiting a stronger binding affinity on naked DNA compared to nucleosomal DNA. This finding is distinct from previous observations where IDRs increase TF binding time on targets ^28,40^. By calculating the target search kinetics with the 3D-diffusion dominant model ^18^, we found that deletion of CTD resulted in a twofold increase in both free diffusion time (τ*_free_*) and target search time (τ*_search_*) between two specific binding events (Fig. 3F and Supplementary Fig. 3H; Supplementary Table 3), suggesting less efficient target search. These findings indicated that while the DBD of FOXA2 determined the DNA binding affinity, its non-DNA binding and histone-interacting CTD promoted its confined diffusion, frequent chromatin sampling, and efficient target search.

### Disordered CTD boosts pioneer activity

The unique kinetics of PTF-chromatin interaction prompts the question of its contribution to pioneer activities, including chromatin binding and chromatin opening. To avoid indirect effects, we expressed Flag-tagged FOXA2 and ΔCTD with the Tet-on system for four days in U2OS cells, and performed ChIP-seq and ATAC-seq (Fig. 4, A and B; Supplementary Fig. 4, A to F; Supplementary Table 4). U2OS cells were chosen as they did not express any of the FOXA family proteins. FOXA2 functioned as a typical PTF, as 86% of its binding sites were initially in closed chromatin and 43% of them were induced to open state (Fig. 4A). In comparison to FOXA2, its ΔCTD mutant showed clear defects in pioneer activity, losing stable binding at 48.7% of FOXA2 sites (Fig. 4B; Supplementary Fig. 4A) and displaying a deficiency in inducing chromatin opening, even at ΔCTD-bound sites (Fig. 4A; Supplementary Fig. 4, F and G). Notably, the impaired binding upon CTD deletion was preferentially in closed chromatin rather than open chromatin (Fig. 4A). These results showed that CTD of FOXA2 greatly enhanced its pioneer activity of binding and opening closed chromatin target sites.

**Figure 4.**
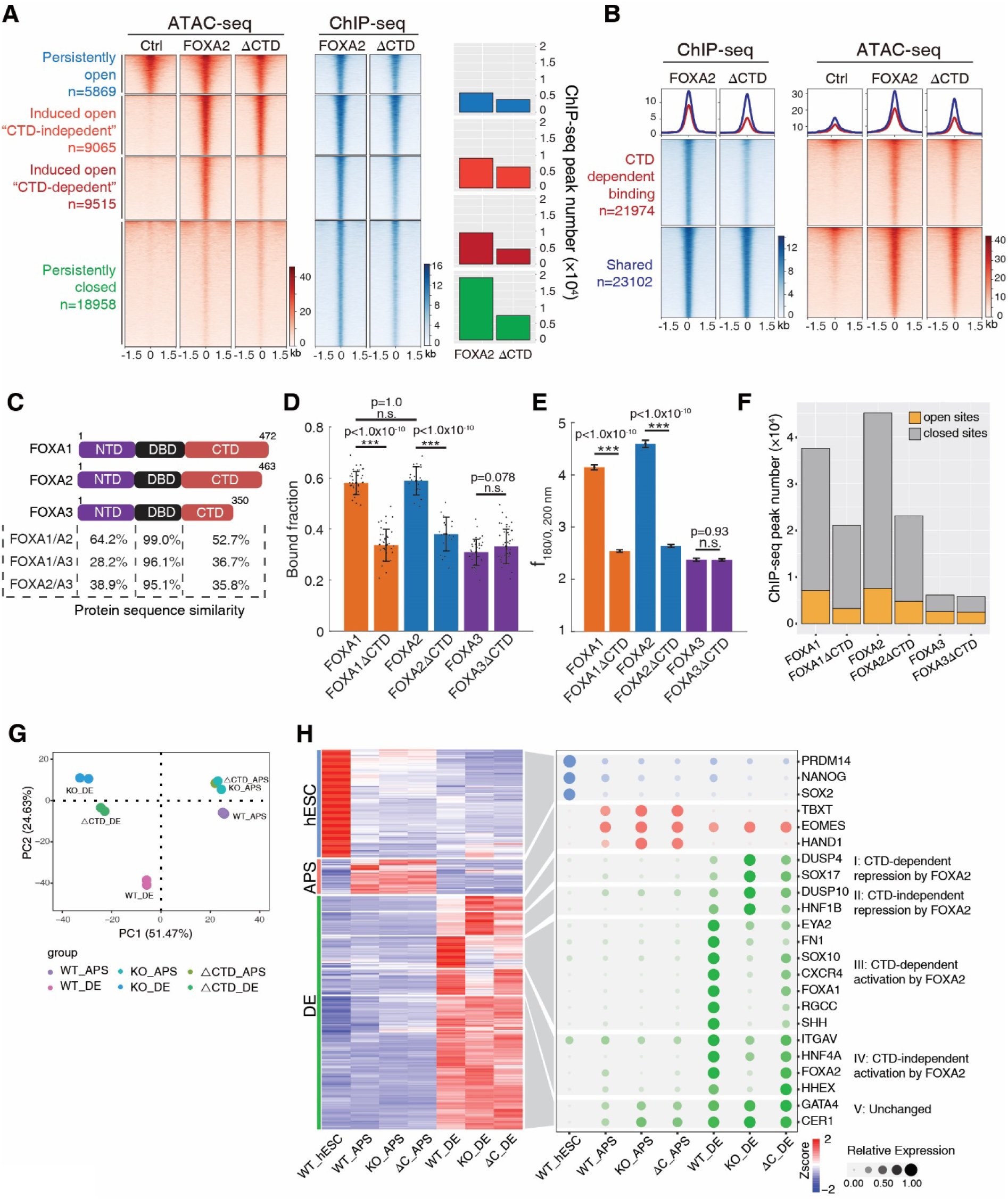
FOXA CTD enhances pioneering closed chromatin and is required for normal DE-specific gene expression. (**A**) Heatmaps of ATAC-seq signal and ChIP-seq signal of indicated samples centered by the FOXA2 binding sites, classified by the change of ATAC-seq signal after four-day expression of FOXA2. The number of ChIP-seq peaks in each group is plotted on the right. (*n* = 2 biological replicates) (**B**) Heatmaps of ChIP-seq and ATAC-seq signal of indicated samples centered by the FOXA2 binding sites, classified by the change of ChIP-seq signal. (**C**) Protein sequence similarity of human FOXA factors. (**D**) Bound fractions (mean ± S.D., dots represent data points) derived from two-state Spot-On analyses of fast spaSMT trajectories for indicated molecules in U2OS cells. (**E**) *f_180/0,_* _200nm_ (peak value of space dependent anisotropy at 200 nm, mean ± SEM) of indicated molecules in U2OS cells. Error bars show standard errors from 50 subsamplings of segment angles with 50% replacement. (**F**) Numbers of ChIP-seq peaks in either open or closed chromatin based on ATAC-seq of wildtype U2OS cells. Shared ChIP-seq peaks of FOXA and FOXA ΔCTD are plotted for the ΔCTD samples. (**G**) Principal component analysis of indicated RNA-seq samples with differentially expressed genes. (**H**) (Left) RNA-seq heatmap of FOXA2 putative targets classified by stage-specific expression in hESC (259 genes), APS (83 genes) and DE (546 genes). (Right) Bubble plot of representative genes in five subgroups of DE genes. (*n* = 2 biological replicates). Significance was calculated using one-way ANOVA followed by Tukey’s multi-comparison, ***p-value<0.001, n.s., not significant. (**D** and **E**) n=33-58 biologically independent cells for each sample, see Supplementary Table 1.

The mammalian FOXA factors, FOXA1, FOXA2, and FOXA3, share well-conserved DBDs but have less conserved CTDs, with CTD_FOXA3_ being the shortest and sharing the least similarity (Fig. 4C). Using fast spaSMT and ChIP-seq, we investigated the chromatin binding of FOXA factors and corresponding ΔCTD variants in U2OS cells (Supplementary Fig. 4, H and I). Our results showed that FOXA3 had a lower chromatin-bound fraction (FOXA1 58.0%, FOXA2 58.9%, FOXA3 30.9%) and weaker space-dependent anisotropic diffusion compared to FOXA1/2 (Fig. 4, D and E). Deletion of CTD_FOXA1_ and CTD_FOXA2_ led to reduced chromatin-bound fractions, while deletion of CTD_FOXA3_ did not. The ChIP-seq profiles of the three FOXA factors were also distinct (Supplementary Fig. 4, J and K). FOXA1/2 bound extensively to closed chromatin sites, whereas FOXA3 had fewer binding sites and tended to bind open chromatin or relatively accessible regions (Fig. 4F and Supplementary Fig. 4L). FOXA1/2 binding to closed chromatin lacking distinguishing histone marks (Supplementary Fig. 4M), consistent with previous observations^12^. Hence, FOXA1/2 appeared to have stronger pioneer activity than FOXA3 at least in this cellular context. Like FOXA2, deletion of CTD_FOXA1_ led to substantial loss of specific FOXA1 binding preferentially in closed chromatin (Fig. 4F). Thus, the disordered CTDs differentiated FOXA factors in their single-molecule confinement dynamics as well as the pioneer capability of binding targets in closed chromatin. These results again emphasized the enhancing roles of the disordered CTDs in chromatin confinement and targeting closed chromatin.

Lastly, FOXA2 plays a pioneering role in the formation and differentiation of definitive endoderm and their derivatives, both in the hESC *in vitro* differentiation model^35^ and in mouse models^32^. Deletion of the histone-interacting region of CTD in mouse FOXA2 impaired embryonic development, altered gene activation and the accessible chromatin landscape^32^. Thus we aimed to examine the roles of FOXA2 and its CTD in hESC differentiation to DE. For this purpose, we generated FOXA2 knockout (KO) hESC clones in addition to ΔCTD clones and conducted RNA-seq analyses on these cells at the hESC, APS, or DE stages (Supplementary Fig. 4, N and O). In examining the gene network transition from hESC to APS and DE, we found that KO cells displayed minor defects in activating APS-specific genes and more apparent defects in activating DE-specific genes (Fig. 4G), and that CTD was required for the normal expression of a subset of them, including SOX17, FOXA1, and CXCR4 (group I and III in Fig. 4H; Supplementary Table 6). These results highlight the importance of CTD for normal DE-specific gene expression.

### Interaction of FOXA2 with mesoscale chromatin domains

We next examined the spatial relationship of FOXA2 with chromatin by super-resolution microscopy. Specifically, we used two-color photoactivated localization microscopy (PALM) to simultaneously detect FOXA2 (tagged with Halo and stained by photoactivatable PA-JF646) and the core histone protein H2B (tagged with photoconvertible fluorescent protein mEosEM^41^) in paraformaldehyde-fixed U2OS cells (Fig. 5A; Supplementary Fig. 5, A to C). Cells with comparable expression level were sorted and selected for imaging experiments. H2B PALM exhibited a higher localization density at the nuclear periphery (Fig. 5A; Supplementary Fig. 5D), where heterochromatin was frequently located, indicating a high degree of chromatin compaction. To quantitatively analyze the cluster sizes of different FOXA2 variants, we used SR-Tesseler^42^, a robust and unbiased segmentation algorithm, and found that full-length FOXA2 formed larger clusters than FOXA2ΔCTD and FOXA2ΔDBD (Supplementary Fig. 5E). Notably, measured by Mander’s correlation, FOXA2 PALM in the same cells showed significantly colocalized distribution with H2B PALM, especially at the nuclear periphery (Fig. 5B and Supplementary Fig. 5F). In contrast, ΔCTD showed less degree of periphery colocalization with H2B and ΔDBD appeared even nuclear distribution. Therefore, the two-color PALM results agreed with genomic results of FOXA2 targeting closed chromatin regions in a CTD-dependent manner.

**Figure 5.**
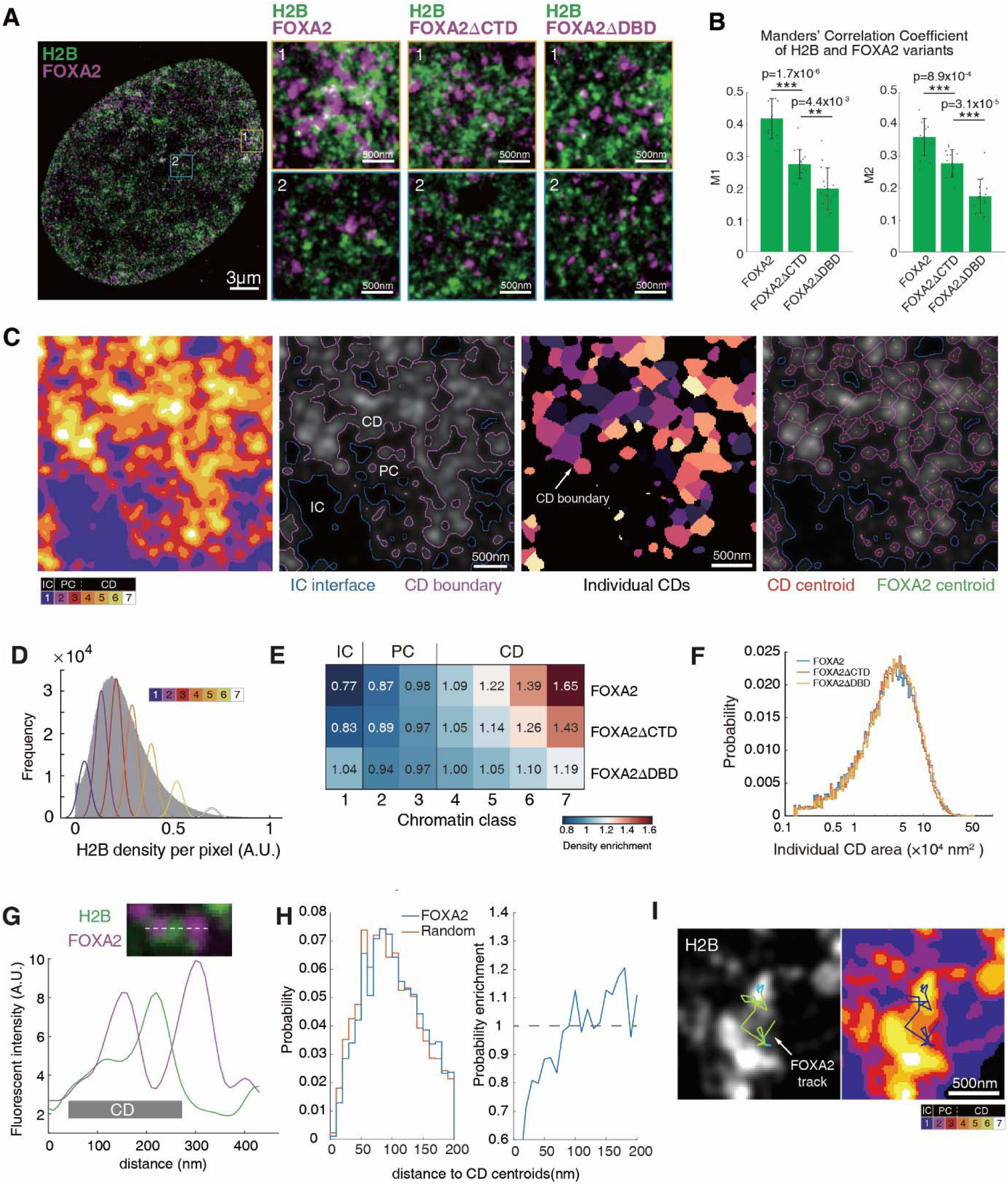
Two-color PALM reveals that FOXA2 targets compacted chromatin domains enriched for nucleosomes. (**A**) Two-color PALM images of fixed U2OS cells that express mEosEM-tagged H2B and Halo-tagged FOXA2 variants labeled with PA-JF646. Blue and yellow boxes indicate representative peripheral and central regions showing the spatial relationship between chromatin and FOXA2 variants. (**B**) Manders’ correlation coefficients (mean ± S.D., dots represent data points) for whole cell show the colocalization of FOXA2 variants and H2B in both directions with auto thresholds. n=12 (FOXA2) or n=14 (FOXA2ΔCTD/FOXA2ΔDBD) biologically independent cells. (**C**) Process of model-based chromatin classification and analysis. (Column 1) Segmentation of chromatin into seven intensity classes based on HMRF model. (Column 2) Blue and magenta contours partition chromatin into three compartments: interchromatin (IC, class 1), perichromatin (PC, class 2-3), and chromatin domain (CD, class 4-7). (Column 3) Seperation of individual chromatin domains. (Column 4) CD centroids (Red) and FOXA2 centroids (Green) relative to chromatin domains. (**D**) Histogram of the H2B intensity of the example nucleus image in grey, overlaid with probability density functions of the seven Gaussians generated by the HMRF model. (**E**) Enrichment or depletion of FOXA2 variants’ density relative to a random distribution for each chromatin class. (**F**) Area distribution of individual CDs. (**G**) Representative image and intensity line plots of a chromatin domain in relation to FOXA2. (**H**) Distribution of distance to the CD centroids from FOXA2 centroids compared to (left) or divided by (right) random localizations. (**I**) Overlaid image of live-cell PALM of H2B and fast spaSMT of FOXA2. The trajectory of FOXA2 is color-coded with green and blue, representing the “free” and “bound” states, respectively. Significance was calculated using one-way ANOVA followed by Tukey’s multi-comparison, ***p-value<0.001, n.s., not significant. (**D** to **H**) n=12 (FOXA2) or n=14 (FOXA2ΔCTD/FOXA2ΔDBD) biologically independent cells.

We further investigated the super-resolved characteristics of FOXA2-bound chromatin with PALM. While nucleosomes displayed discrete nanodomains with a width of a few tens of nanometers (Fig. 5A)^43^, we hypothesized that observed chromatin confinement of FOXA2 was related to chromatin structure at a larger physical scale, as the confined diffusion zones had an averaged radius of ∼200 nm (Fig. 2C). Thus, we employed the hidden Markov random field model (HMRF) to classify chromatin in the nucleus into seven compaction levels and subsequently into three functional compartments representing different chromatin activities based on previous characterizations^44^ (Fig. 5, C and D; Supplementary Fig. 5G). The resulting chromatin compaction landscape featured chains of mesoscale chromatin domain regions (CD) having high nucleosome density (class 4-7), broader perichromatin regions (PC) at the outer rim of CD having lower nucleosome density (class 2-3), and interchromatin regions (IC) depleted of nucleosomes (class 1). By measuring FOXA2 molecule localizations or cluster centroids in chromatin regions of different compaction levels, we found that FOXA2 could bind and even preferentially bound the compacted closed chromatin regions (Fig. 5E; Supplementary Fig. 5, H and I). In comparison, the ΔCTD showed less density enrichment in CDs, consistent with its impaired pioneer activity.

We further subdivided CD regions into single CDs and identified their centroids with the strongest nucleosome intensity. Each cell at the focal plane had ∼ 3000 CDs (Supplementary Fig. 5J) with an average area of 4.4×10^4^ nm^2^ and diameter of 240 nm (Fig. 5F), which was similar to the physical size of confinement zones (Fig. 2C). The coincidence suggested that organization of individual CDs might underlie observed chromatin confinement of FOXA2, in consideration of the tight correlation among chromatin confinement, chromatin-bound fraction and successful binding at closed chromatin sites. By quantifying the distances between the CD centroids and FOXA2 centroids, we showed that FOXA2 spanned across the entire CD entity with a slight preference to the periphery (Fig. 5, G and H). Notably, we observed the confinement trajectories of single FOXA2 molecules within and surrounding the CDs by combining live-cell PALM and fast spaSMT techniques (Fig. 5I, Supplementary Fig. 5K). Moreover, apart from using model-based chromatin segmentation, we have also taken a parallel analysis approach by dividing the nucleus into nine equal-area regions with increasing H2B density and obtained consistent conclusions (Supplementary Fig. 5, L and M). Together, these results provided super-resolution imaging evidence that FOXA2 targeted mesoscale compacted chromatin domains and that CTD facilitated this process. Importantly, our findings suggested that interaction of FOXA2 with mesoscale chromatin domains might underlie the diffusion confinement within a characteristic space and result in efficient pioneer activity.

### Kinetic advantage of confined target search

We next investigated how PTFs might gain kinetic advantages from confined diffusion in binding energetically unfavorable and nucleosome-embedded targets. In order to simulate the confined target search process, we conducted coarse-grained computer simulations of chromatin as a chain of monomers, partially containing a cognate binding site (CBS) of the TF ^38,45^. The simulations assigned a monomer radius of 200 nm to model the spatial volume of transient chromatin confinement observed in experiments (Fig. 6A; Supplementary Fig. 6, A; Supplementary Table 7; Supplementary Video 6). The TF underwent Brownian motion in the nuclear space between chromatin monomers (free nuclear diffusion), and once it entered a monomer, it diffused inside at a lower diffusion coefficient and was confined with a certain probability to exit (P_exit_ < 1) (confined diffusion). While diffusing inside the chromatin monomers, the TF could pause briefly at a given rate to model non-specific binding and have chances to contact the CBS within.

**Figure 6.**
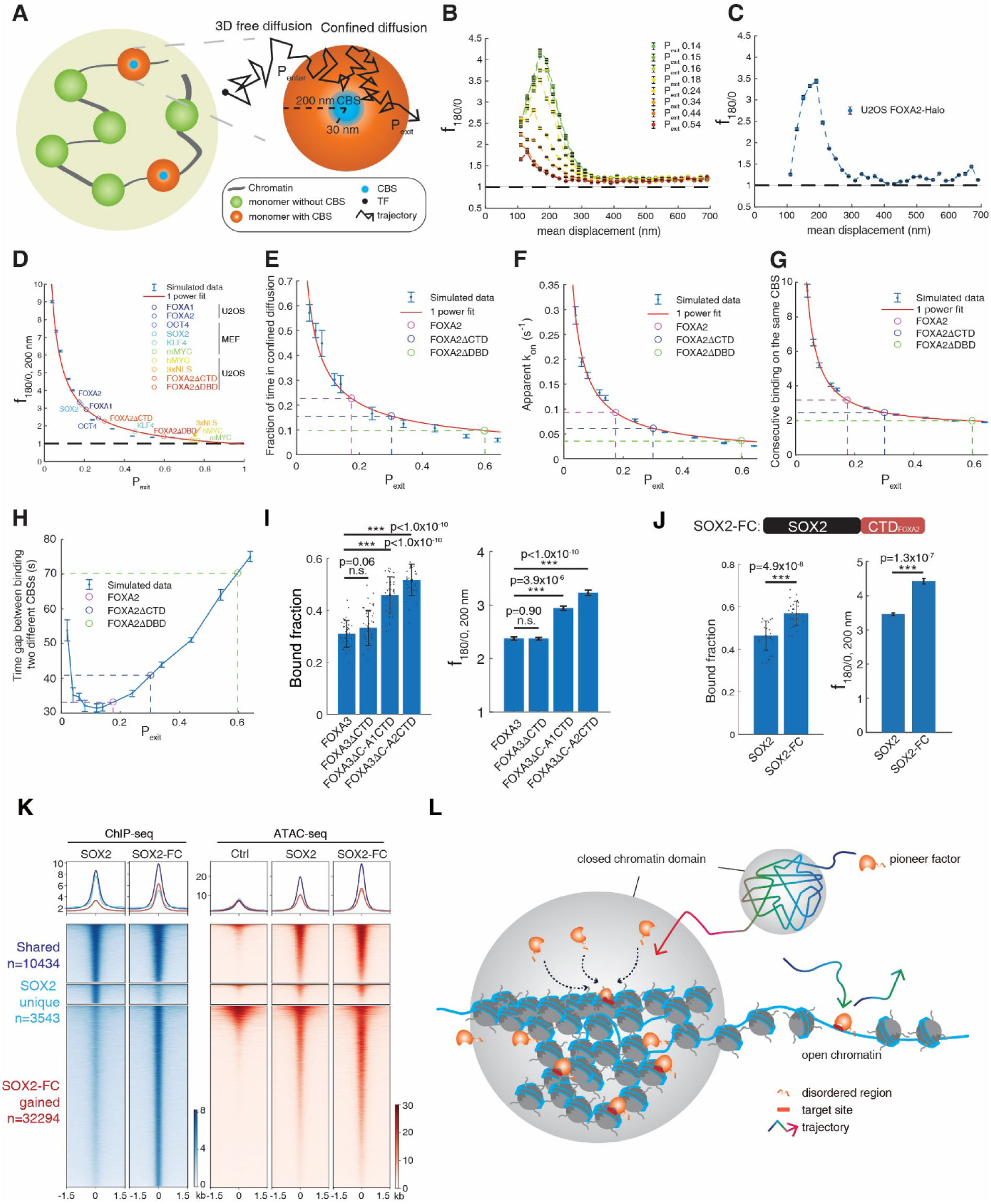
Computer simulations reveals the kinetic advantage of confined target search. (**A**) Chromatin is modeled as a chain of monomers with or without cognate binding site (CBS). TF undergoes 3D free diffusion outside of the monomers, and confined diffusion within the monomers with certain probabilities to enter (P_enter_) and exit (P_exit_). (**B** and **C**) Space-dependent anisotropy examined by *f_180/0_*against mean displacement length of selected free segments, using simulation-generated trajectories with variable P_exit_ (**B**) or experimental data of FOXA2 (**C**). (**D**) The plot of space-dependent anisotropy, measured by *f_180/0,_*_200nm_ (mean ± SEM), as a function of P_exit_. The plot was fitted by one-power model, and the P_exit_ of indicated molecules was extrapolated using experimental *f_180/0,_* _200nm_ values. (**E** to **G**) Indicated kinetics calculated using simulated trajectories at various P_exit_. The simulated data was well fitted by one-power model, and the corresponding y-axis values of indicated molecules were calculated with extrapolated P_exit_ from (**D**). n=12 independent simulations. (**H**) Time gap between binding two different targets (two CBSs in different monomers) calculated using the simulated trajectories at various P_exit_, overlaid with estimated values of indicated molecules using the extrapolated P_exit_. n=12 independent simulations in (**B, D** to **H**) For each round of simulations, there are 5,000 diffusion steps per round (**B, D** and **E**) or 500 times of CBS binding per round (**F-H**). Data are presented as mean ± S.D. in (**E** to **H**). (**I** and **J**) Bound fractions (mean ± S.D., dots represent data points) and *f_180/0,_* _200nm_ (mean ± SEM) derived from fast spaSMT trajectories for indicated molecules in U2OS cells. FOXA3ΔC-A1CTD and FOXA3ΔC-A2CTD stand for fusion protein of FOXA3ΔCTD with the CTD of FOXA1 and FOXA2 respectively. SOX2-FC stands for fusion protein of SOX2 and CTD of FOXA2. (**K**) The heatmaps and averaged profiles of ChIP-seq (left) and ATAC-seq (right) of U2OS cells expressing indicated proteins. Ctrl stands for wildtype U2OS cells. (*n* = 2 biological replicates) (**L**) The working model of PTF target search. Significance calculated using one-way ANOVA followed by Tukey’s multi-comparison in (**I**) and two-sided t-test in (**J**), *p-value<0.05, **p-value<0.01, ***p-value<0.001, n.s., not significant. (**I** and **J**) n=23-36 biologically independent cells for each sample, see Supplementary Table 1.

Synthetic TF trajectories were generated by assigning variable P_exit_ to model different levels of confinement, with the lowest P_exit_ representing the strongest confinement. The space-dependent diffusion anisotropy from synthetic TF trajectories showed striking resemblance to those from experimental data of TFs, peaking around 200 nm (Fig. 6, B and C). The pattern of space-dependent anisotropy remained with variable virtual frame rate (Supplementary Fig. 6B), and showed inverse correlation with P_exit_ (Fig. 6D), confirming that space-dependent trajectory anisotropy measured diffusion confinement. With increasing level of confinement, the target search mechanism of TFs gradually switched from free diffusion dominant to confined diffusion dominant (Fig. 6E), resulting in higher target search efficiency (Fig. 6F) and repetitive trials of target binding (Fig. 6G) which might be essential for PTF to disturb local nucleosome compaction and increase the chance of binding partially exposed motif on nucleosomes. Using *f*_180/0, 200nm_ measured from experiments, we extrapolated the corresponding P_exit_ of TFs and calculated their target search kinetics (Fig. 6D; Supplementary Table 8). The pioneer factors FOXA1, FOXA2, and SOX2 showed the strongest diffusion confinement, with a predicted P_exit_ of FOXA2 at 0.17 and a fraction of confined diffusion at 23% (Fig. 6E). In contrast, the predicted P_exit_ values of MYC, NLS and FOXA2 ΔDBD were close to 1, indicating little confinement. Compared to FOXA2, its ΔCTD mutant showed a higher predicted P_exit_ at 0.3, a lower fraction of confined diffusion at 15%, a lower On-rate to CBS, and less repetitive binding on CBS. The changes in target search kinetics may explain its impaired pioneer activity in binding targets in closed chromatin.

We speculated that extremely strong confinement (a very small P_exit_) could trap TFs unproductively within monomers without CBS for extended periods, negatively impacting the overall efficiency of target search throughout the nucleus. Therefore, we calculated the search time between binding two different CBSs in separate monomers and found that extremely small or large P_exit_ values indeed resulted in long search time (Fig. 6H). Importantly, we identified an optimal P_exit_ that minimized the search time, thereby maximizing efficiency. Notably, the predicted P_exit_ values of FOXA1, FOXA2, and SOX2 fell around the optimal P_exit_. These results suggested pioneer factors may have evolved their protein-nucleosome interaction to achieve an optimal degree of confinement for efficient target searching without becoming overly trapped.

Additionally, we explored an alternative model wherein the observed confinement could be attributed to repetitive binding to specific target sites that are clustered. To test this hypothesis, we modified the chromatin simulation by placing multiple binding sites (5, 50, or 100 CBSs) clustered within a single 200-nm radius chromatin monomer (Supplementary Fig. 6A). Subsequently, we performed the diffusion simulation without confinement by setting P_exit_ to 1 and examined the anisotropy of the diffusion fraction in simulated molecule trajectories as described earlier. The results revealed an absence of spatial confinement of the diffusing molecules (Supplementary Fig. 6C), despite the default clustering of bound molecules in this case. Therefore, it became evident that clustered target sites could not explain the observed anisotropy of diffusing PTFs.

### Modulation of target search by non-DNA binding regions

Our results have established the essential roles that the non-DNA binding disordered regions could play in target search dynamics and pioneer activity. Thus, we speculated that such activity could be used to alter the pioneer activity of other TFs through protein fusion. Indeed, fusion of CTD_FOXA1_ or CTD_FOXA2_ to FOXA3ΔCTD greatly increased the chromatin-bound fraction (from 33.2% to 45.8% and 51.7%) and space-dependent anisotropy (*f*_180/0, 200nm_ from 2.37 to 2.94 and 3.23) as measured by fast spaSMT (Fig. 6I). This finding indicated that CTD_FOXA1/2_ could increase the chromatin interaction and confined diffusion of a homologous DNA binding protein. To further test whether this strategy can be applied to TFs from a different family, we used the Tet-on system to express SOX2 and a fusion protein of SOX2 and CTD_FOXA2_ (SOX2-FC) in U2OS cells (Supplementary Fig. 6D). By fast spaSMT in living cells, SOX2-FC showed a higher chromatin-bound fraction and higher diffusion anisotropy than SOX2 (Fig. 6J), indicating that addition of CTD further enhanced chromatin interaction and chromatin confinement. We further examined them using ChIP-seq and ATAC-seq.

Notably, while SOX2 functioned as a typical PTF, binding targets that were originally in a closed state and increasing their chromatin accessibility, SOX2-FC exhibited remarkably enhanced pioneer activity. This was evident in its ability to bind significantly more SOX2 target sites in closed chromatin and result in a higher level of chromatin opening (Fig. 6K and Supplementary Fig. 6, E and F). Importantly, the gained binding sites of SOX2-FC were *bona fide* SOX2 targets that contained SOX2 consensus motif and shared little overlap with FOXA2 binding sites (Supplementary Fig. 6G). These results highlighted a significant enhancement of SOX2’s pioneer activity through fusion with FC. Thus, a non-DNA binding disordered region could promote TF targeting of closed chromatin, offering a method to improve pioneer activity by altering the target searching process.

## Discussion

Here we revealed a previously unrecognized “confined target search” mechanism employed by PTFs to target compacted closed chromatin (Fig. 6L). Unlike the previously identified 3D diffusion dominant model, PTFs alternate between non-exhaustive 3D free diffusion in the nucleus and confined diffusion in mesoscale compacted chromatin domains. This reduces target search time and enables PTFs to repetitively probe nucleosomal targets, thereby enhancing pioneer activity in closed chromatin. Importantly, this process depends on the non-DNA binding disordered regions of PTFs, facilitating confined diffusion through flexible and frequent interactions with core histones enriched in closed chromatin. Our findings also provides insights for identifying new natural PTFs and engineering synthetic ones.

Our results underscore the crucial role of chromatin spatial organization in PTF target search. While the epigenetic determinants guiding PTF binding to nucleosomal targets remain unclear^12^, our super-resolution imaging revealed the presence of spatially organized mesoscale compacted chromatin domains, acting as landing pads that attract PTFs through interactions with their disordered regions. Subsequent transient diffusion confinement within these chromatin domains further reduces the search space required to find targets. Additionally, it increases the frequency of binding attempts by PTFs, potentially alleviating local nucleosome compaction and transiently exposing buried motifs for stable PTF binding ^46,47^. This mechanism aligns with prior observations including slow nuclear mobility of FOXA factors^25,26^, non-sepecific chromatin sampling of FOXA2 ^12^, Zelda hubs^27^ and OCT4 interactions with histones^47^, suggesting a shared mechanism among PTFs in leveraging chromatin organization for target search in closed chromatin. Hence, our study unveils a remarkable mechanism through which PTFs exploit chromatin spatial architecture to access closed chromatin.

Our findings highlight the essential role of disordered regions in pioneer function. The nucleosomal binding capability of PTFs is generally thought to be encoded within the well-structured DBD, and the fine structural features of PTF-nucleosome interactions have been characterized ^2,48^. As the ternary complex of TF-bound nucleosome is relatively unstable, a TF with strong DNA affinity could likely stabilize its binding to a partially dissociated nucleosome. Consistently, our results revealed that FOXA2 remained bound to targets in living cells for a relatively long period of time. This observation is consistent with the extended residence time of *Drosophila* GAGA factor which is also a PTF ^28^. However, the DBD alone may not guarantee pioneer activity, as TFs from the same family with highly similar DBDs do not always function as PTFs; in line with this notion, FOXA1/2/3 showed distinct levels of capacity to bind closed chromatin in the same cellular context. TFs commonly contain IDRs that mediate multivalent interactions and facilitate a wide variety of biological activities, including their binding dynamics ^40,49–52^. The potential contribution of diverse disordered regions to pioneer activity has only recently begun to be unraveled ^3,53,28,54,47^. In this study, we found that the IDR of FOXA2 played a surprisingly important role in the dynamic process of target search, particularly for closed targets. While it promoted target association rate, its deletion did not seem to affect dissociation rate. This is different from the IDR of GAGA factor ^28^, suggesting it is unlikely all pioneer factors use exactly the same mechanism. Moreover, the structured DBD and the disordered non-DBD regions could function as modules, contributing to stable target binding and target search in closed chromatin, respectively, although one indeed influences the other. Given that PTFs are often mutated in both DBD and non-DBD regions across various cancer types, comprehending the fundamental properties of PTFs holds significant value in human cancer research. Our study encourages further exploration of the diverse disordered regions within other PTFs.

## Acknowledgements

We are grateful to Drs. Robert Tjian, Xavier Darzacq, and their laboratory members for experiment assistance and discussions in initiating this study. We thank Rachel Salomon and Gina Dailey for assistance in cell culture and cloning. We thank Dr. Assaf Amitai for providing computer code and discussion on kinetic simulation of target search. We thank Dr. Pingyong Xu for sharing the mEosEM plasmid. We thank the National Center for Protein Sciences at Peking University for providing FACS service and confocal microscopes. We thank the following for funding support: National Key Research and Development Program of China 2020YFA0509502 (WD); National Natural Science Foundation of China 32170566 (WD); Beijing Advanced Innovation Center for Genomics at Peking University (WD); Peking-Tsinghua Center for Life Sciences (WD); National Natural Science Foundation of China T2225001 (HG).

## Author Contributions Statement

Design of experiments: WD, ZW, BW and DN; Live-cell SMT and analysis: ZW and WD; ChIP-seq and ATAC-seq: DN; RNA-seq: WD and CC; Genomics analysis: BW, DN and YB; Two-color PALM and analysis: BW; Computer simulation of target search: HG, ZW and BW; Lentivirus production and MEF SMT: YC; hESC culture and differentiation: WD, YC and KML; Cell line construction: WD, ZW, DN, BW and YB; Halo dye synthesis: LDL; Writing – original draft: WD, ZW, BW and DN; Writing – review and editing: CC, KML and HG.

## Competing Interests Statement

The authors declare no competing interests.

---

**Supplementary Fig. 1.**
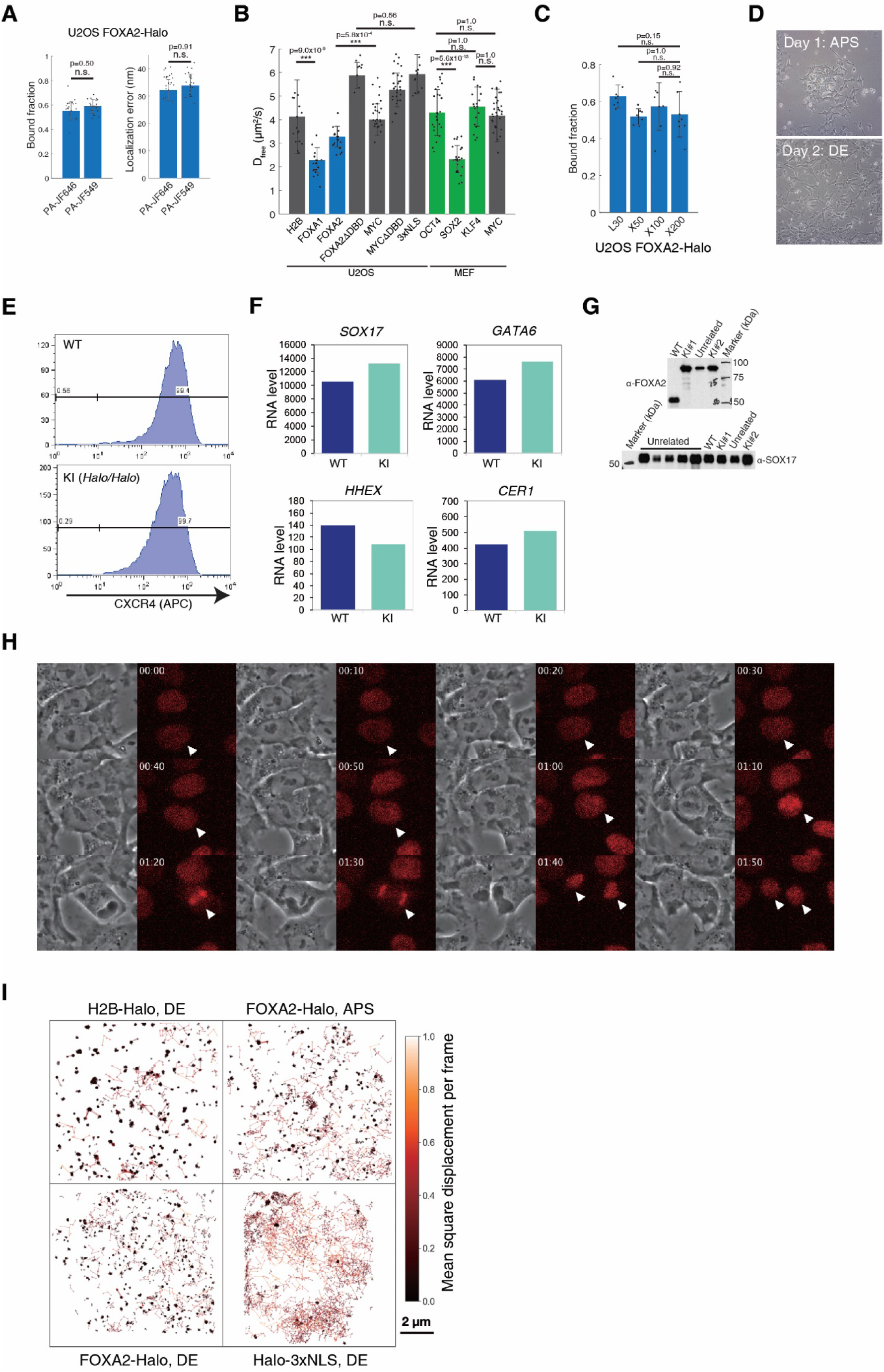
Fast spaSMT and hESC knock-in clones. (**A**) Two PA dyes generated comparable results in fast spaSMT experiments. Bound fraction (mean ± S.D.) (left) and localization error (mean ± S.D., dots represent data points). (right) were derived from two-state Spot-On analyses of fast spaSMT trajectories of FOXA2-Halo in U2OS cells. n=21 biologically independent cells. (**B**) Diffusion coefficient of free molecules (mean ± S.D., dots represent data points) derived from two-state Spot-On analyses of fast spaSMT trajectories of indicated molecules in U2OS cells or MEF cells. n=16-58 biologically independent cells for each sample, see Supplementary Table 1. (**C**) Bound fraction (mean ± S.D., dots represent data points) of FOXA2-Halo that were exogenously expressed in U2OS cells with the low-expression L30 promoter or by Tet-on system with doxycycline induction 50 ng/mL (X50), 100 ng/mL (X100) and 1μg/mL (X1000). n=8 biologically independent cells. (**D**) Cell morphology of hESCs differentiated APS cells (left) and DE cells (right). (**E** to **H**) The knock-in FOXA2-Halo/Halo homozygous hESC clones (KI) had the same differentiation potential to DE as the WT hESCs, as shown by flow cytometry with DE-specific surface marker CXCR4 (**E**), RT-qPCR of DE-specific genes (n=1 biological replicate) (**F**), western blots of DE lysates from WT and two KI clones with indicated antibodies (**G**), and time-lapsed phase-contrast and fluorescence images (**H**) showing mitotic chromosome binding. The time stamp format is “hour:min”. (**I**) Randomly selected 1000 fast spaSMT trajectories of indicated molecules were plotted with color-coded mean square displacement per frame. Significance was calculated using two-sided t-test in (**A**), one-way ANOVA followed by Tukey’s multi-comparison in (**B** and **C**), *p-value<0.05, **p-value<0.01, ***p-value<0.001, n.s., not significant.

**Supplementary Fig. 2.**
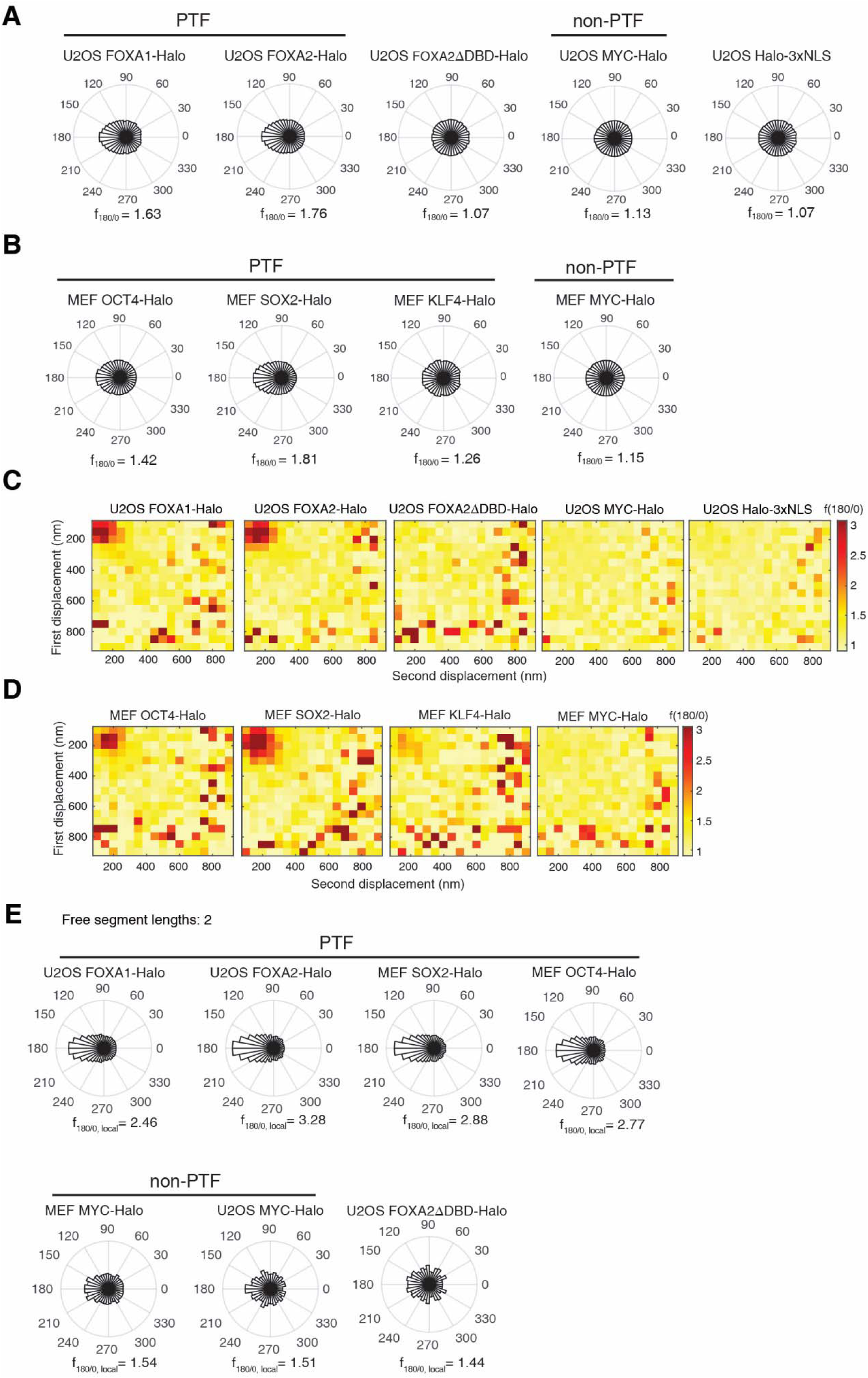
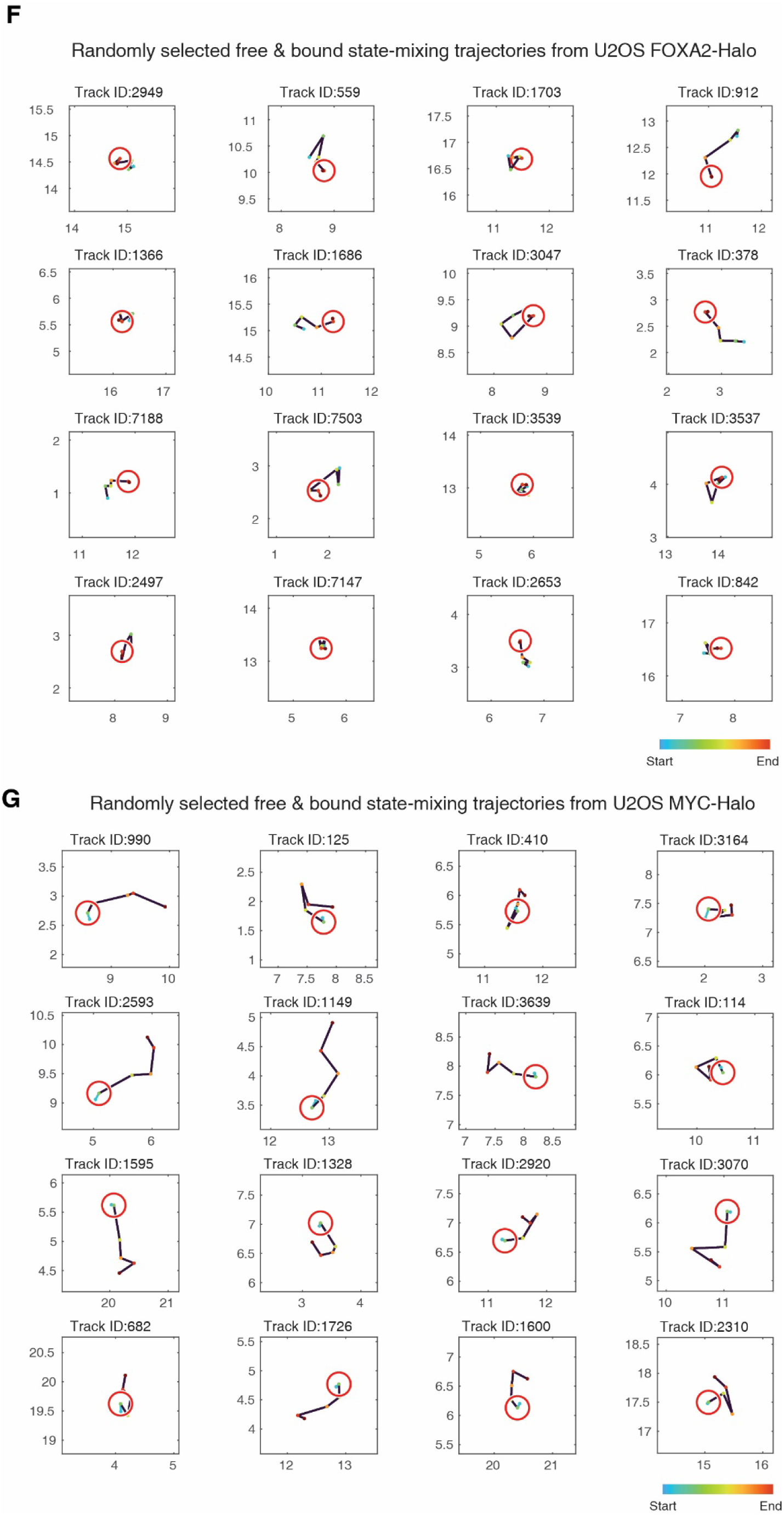
Anisotropic diffusion of PTFs. (**A** and **B**) The angle distribution histograms and *f_180/0_* values of indicated molecules in (**A**) U2OS cells and (**B**) MEF cells. (**C** and **D**) Anisotropy heatmaps showing *f_180/0_* versus length of displacements for indicated molecules in (**C**) U2OS cells and (**D**) MEF cells. (**E**) The angle distribution histograms and *f_180/0,local_*calculated from two segments adjacent to a binding event (local anisotropy). (**F** and **G**) Randomly selected free and bound state-transition trajectories of exogenously expressed (F) FOXA2-Halo and (G) MYC-Halo in U2OS cells. Red circles of 200 nm radius are drawn centering locations of the “bound” events. The free segments are colored black, while locations are color-coded by temporal order. (**A**-**E**) Results are from n=18-58 biologically independent cells for each sample, see Supplementary Table 1.

**Supplementary Fig. 3.**
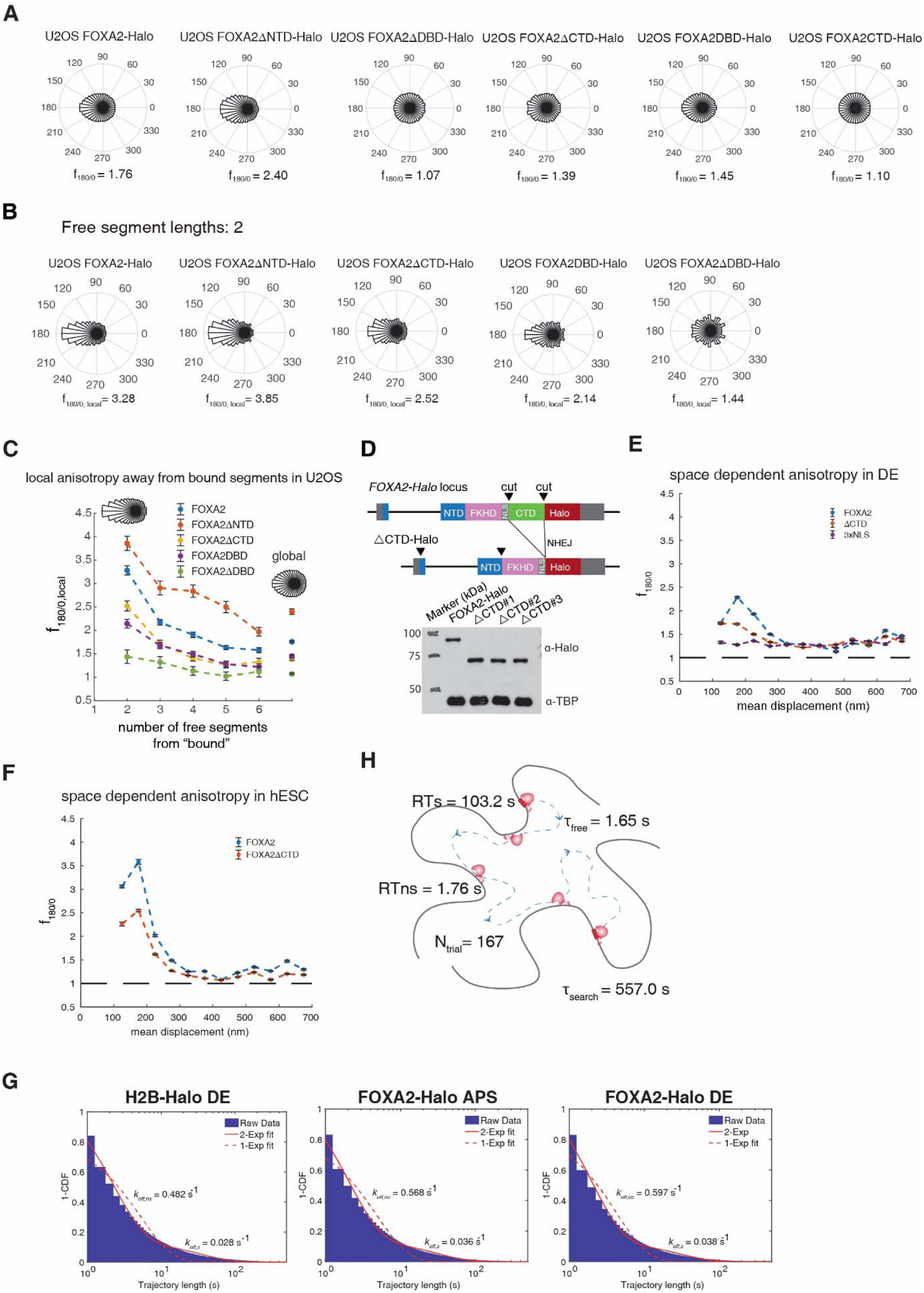
CTD-dependent confined target search of FOXA2 is mediated by CTD-nucleosome interaction. (**A**) The angle distribution histograms and *f_180/0_*values of indicated molecules in U2OS cells. (**B**) The angle distribution histograms and *f_180/0, local_* calculated from two segments adjacent to a binding event. (**C**) *f_180/0, local_* (mean ± S.D.) were calculated from a variable number of free segments before and after a binding event (local anisotropy, left) and compared to *f_180/0_* of all free segments (global anisotropy, right). The representative angle distribution histograms of FOXA2 are shown on the top. (**D**) Illustration of CRISPR/Cas9 mediated non-homologous end joining (NHEJ) to generate hESC clones deleting the CTD (ΔCTD-Halo) based on the FOXA2-Halo KI hESC clone. Western blots of DE lysates from indicated hESCs with indicated antibodies. *, FOXA2-Halo; **, △CTD-Halo. (**E** and **F**) Space-dependent anisotropy examined by *f_180/0_*against mean displacement length of selected free segments, calculated from fast spaSMT of indicated molecules in (E) DE cells and (F) hESC cells. (**G**) Linear-log survival curves of slowSMT trajectories of the exogenously expressed H2B-Halo and endogenously tagged FOXA2 or ΔCTD in DE cells. (**H**) Calculated target search kinetics of FOXA2 between two specific targets interspersed with 3D free diffusion and non-specific bindings. Grey line: chromatin. Red protein: FOXA2. (**A-C** and **E-G**) Results are from n=12-58 biologically independent cells for each sample, see Supplementarty Table 1.

**Supplementary Fig. 4.**
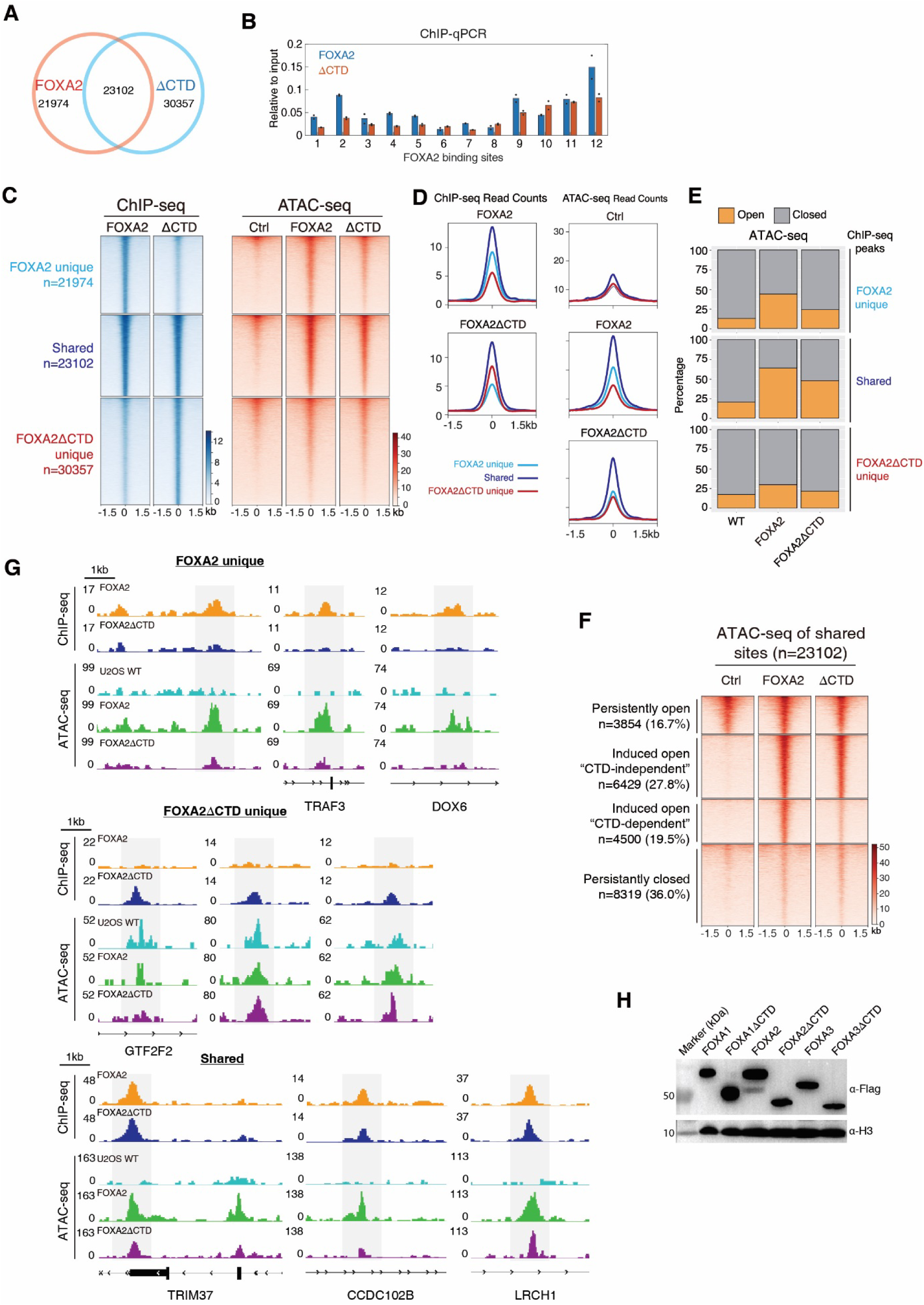
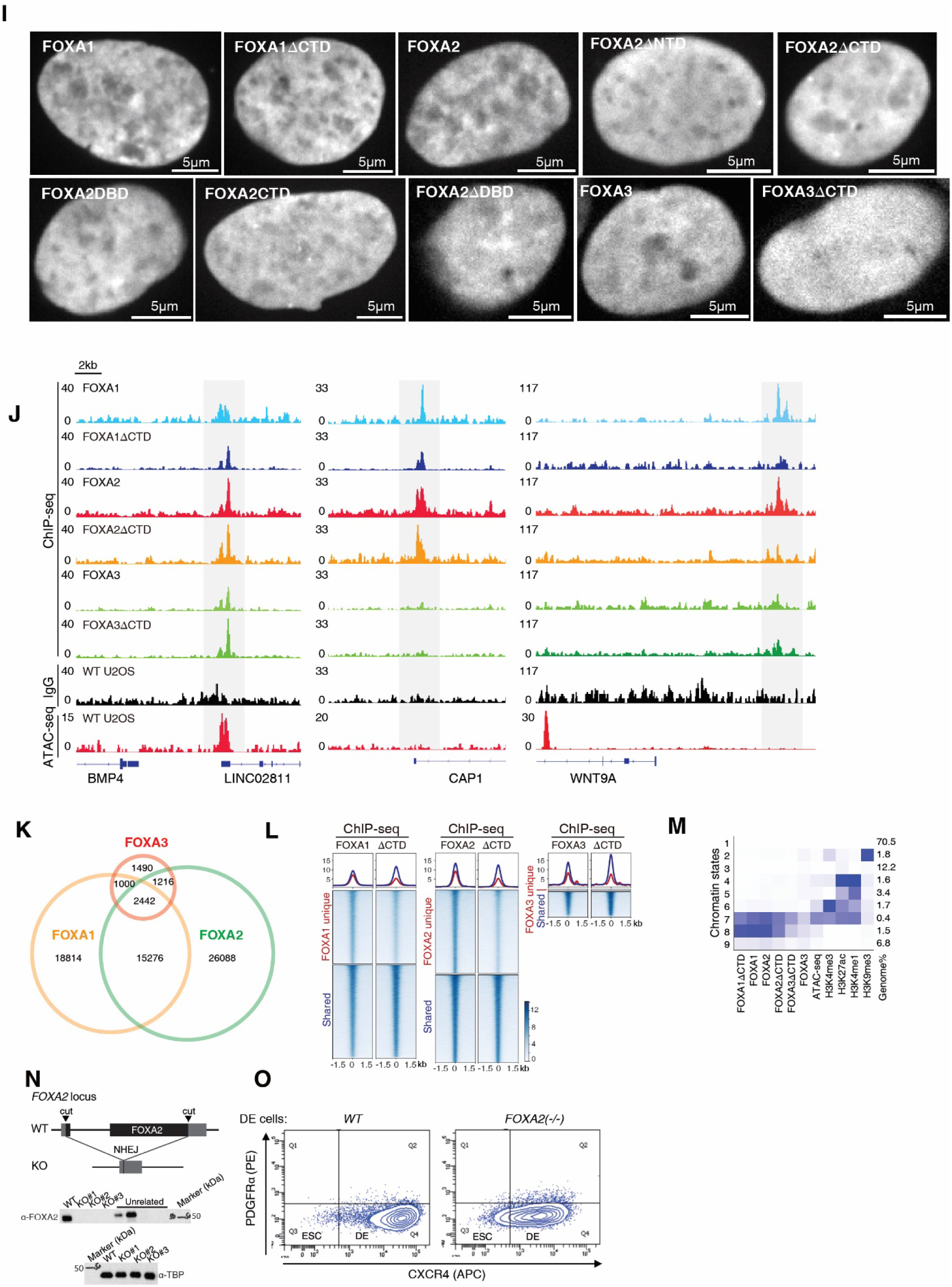
Functional roles of CTD in FOXA pioneer activity and in DE formation. (**A-G**) Analysis of Flag ChIP-seq and ATAC-seq of U2OS cells with exogenous expression of Flag-tagged FOXA2 or ΔCTD. (**A**) Venn diagrams of ChIP-seq peaks from indicated samples. (**B**) ChIP-qPCR results of randomly selected 12 FOXA2 binding sites (*n* = 2 biological replicates, dots represent data points, bar height represents the mean). Heatmaps (**C**) and averaged profiles (**D**) of ChIP-seq and ATAC-seq of indicated samples. (**E**) Percentage of ChIP-seq peaks in open and closed states. (**F**) Heatmap of ATAC-seq at the shared sites by FOXA2 and ΔCTD, where the sites were further classified using the change of ATAC-seq signal and CTD dependency. (**G**) ChIP-seq and ATAC-seq tracks of representative chromatin regions in each cluster. (**H-M**) Analysis of Flag ChIP-seq of U2OS cells with exogenous expression of Flag-tagged FOXA1/2/3 and their ΔCTD mutants. (**H**) Western blots of the whole cell (repeated twice independently with similar results) and (**I**) Confocal images of the indicated cell lines showed comparable expression levels (repeated twice independently with similar results). (**J**) ChIP-seq and ATAC-seq tracks of representative chromatin regions. (**K**) Venn plots of FOXA ChIP-seq peaks in U2OS cells.(**L**) ChIP-seq heatmaps of FOXA1/2/3 and their ΔCTD mutants at FOXA1/2/3 ChIP-seq peaks. (**M**) ChromHMM using epigenetic markers, genome-wide binding profiles of transcription factor and chromatin accessibility. The blue shading depicts the average intensity of a particular mark across each chromatin state. (**N-O**) hESC RNA-seq analysis. (**N**) Illustration of CRISPR/Cas9 mediated knock-out (KO) and western blots of DE lysates (repeated twice independently with similar results). (**O**) Flow cytometry of indicated DE cells with DE surface markers (CXCR4^high^, PDGFRl1l^neg^).

**Supplementary Fig. 5.**
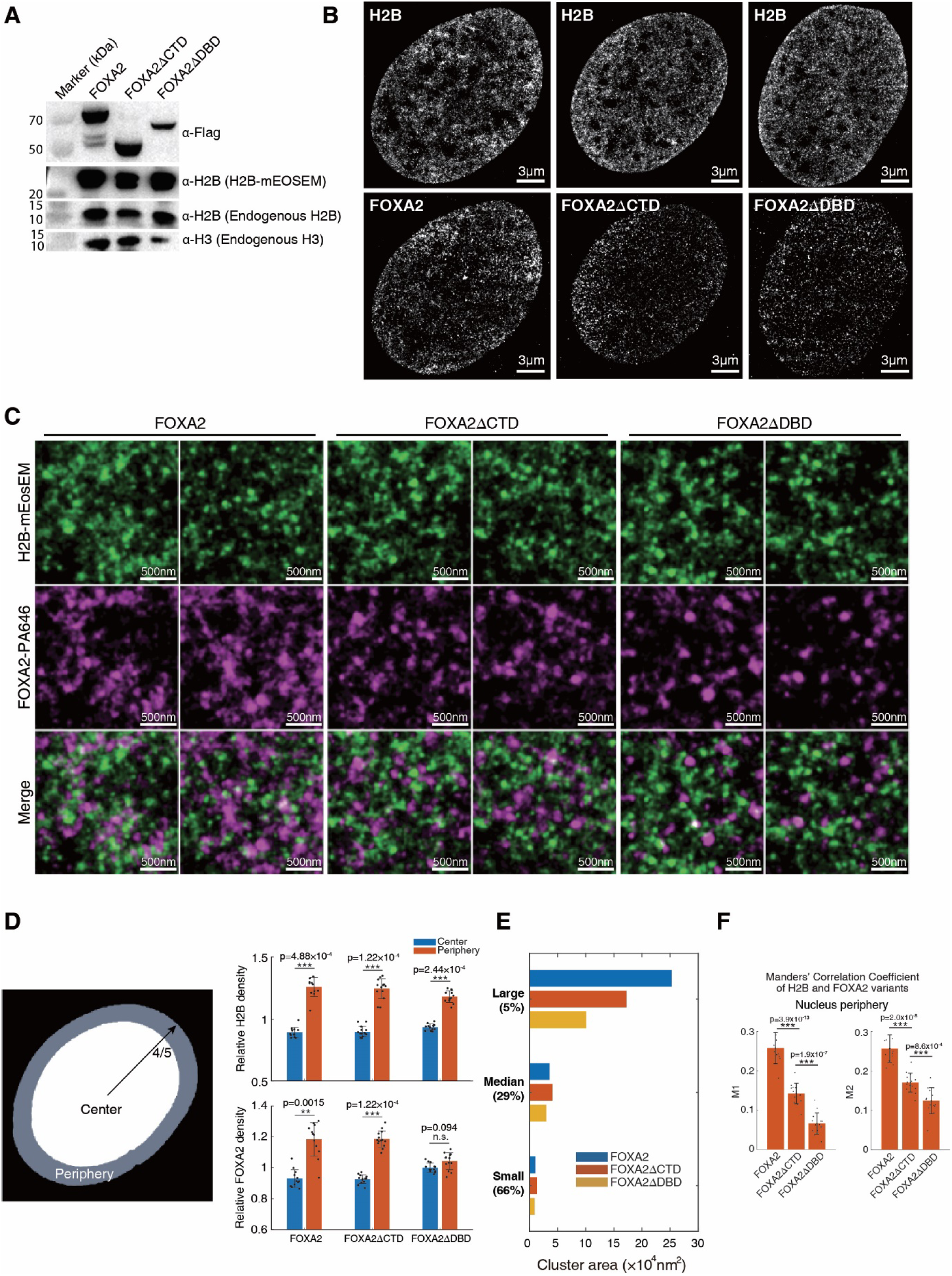
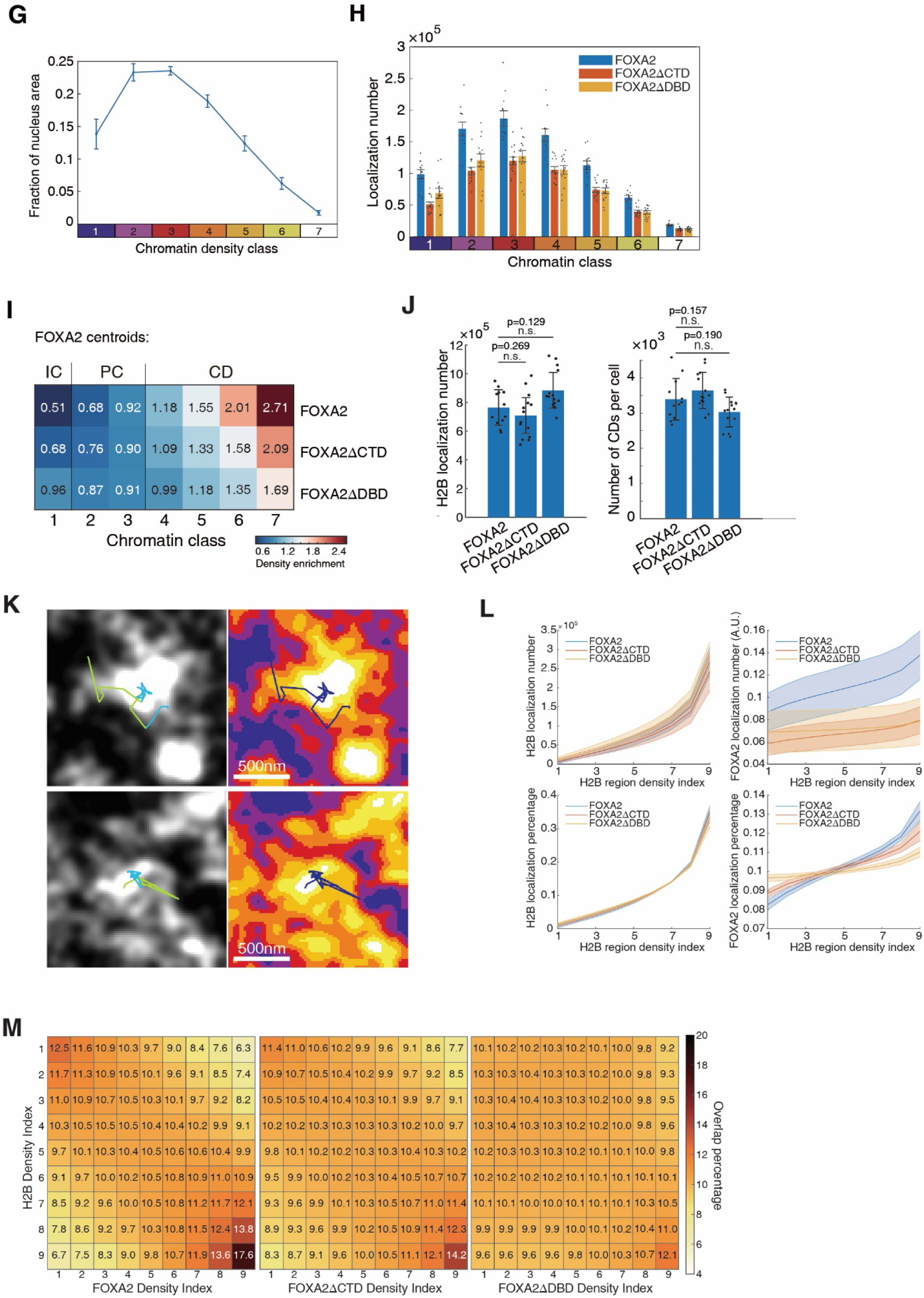
Two-color PALM of chromatin and FOXA2. (**A**) Western blots of the whole cell lysates from the indicated cell lines with indicated antibodies. **(B**) Two-color PALM images of fixed U2OS cells exogenously expressing mEosEM-tagged H2B and Halo-tagged FOXA2 variants labeled with PA-JF646. **(C)** Additional representative two-color PALM images. **(D)** (Left) The cell nucleus was split at four fifth of radius into “center” and “periphery”. (Right) Relative localization density (mean ± S.D., dots represent data points) of H2B and FOXA2 variants in nucleus center and periphery. (**E**) Bar plot displays the cluster areas of FOXA2 variants grouped into small, median, and large sizes based on percentiles. The classification is as follows: 0-66% as small, 66-95% as median, and 95-100% as large. (**F**) Manders’ correlation coefficients (M1 and M2) (mean ± S.D., dots represent data points) of FOXA2 variants and H2B in both directions using signals from the nucleus periphery. **(G)** Fractions (mean ± S.D.) of nucleus area occupied by the indicated chromatin class in FOXA2. (**H**) PALM localization numbers (mean ± SEM, dots represent data points) of FOXA2 variants in different chromatin classes. (**I**) Density enrichment of FOXA2 variants’ centroids relative to a random distribution in each chromatin class. (**J**) (Left) Localization number (mean ± S.D., dots represent data points) of H2B-mEosEM per cell. (Right) Number of separated CDs per cell (mean ± S.D., dots represent data points). (**K**) Overlaid image of live-cell PALM of H2B and fast spaSMT of FOXA2.The trajectory of FOXA2 is color-coded with green and blue, representing the “free” and “bound” states, respectively. (**L**) The nucleus was divided into nine equal-area regions with increasing H2B density. (Top) The localization numbers (mean ± S.D.) of H2B and FOXA2 variants in each region. (Bottom) The localization percentage (mean ± S.D.) of H2B and FOXA2 variants in each region. (**M**) The nucleus was divided into nine equal-area regions with either increasing H2B density or increasing FOXA2 density. Heatmap showing area overlap between the two in the form of percentage. Error bars indicate the standard deviation. (**D, F, G, H, J**) n=12 (FOXA2) or n=14 (FOXA2ΔCTD/FOXA2ΔDBD) biologically independent cells. Significance was calculated using Wilcoxon signed-rank test in (**D**), one-way ANOVA followed by Tukey’s multi-comparison in (**F**), or Wilcoxon rank-sum test in (**J**), **p-value<0.01, ***p-value<0.001, n.s., not significant.

**Supplementary Fig. 6.**
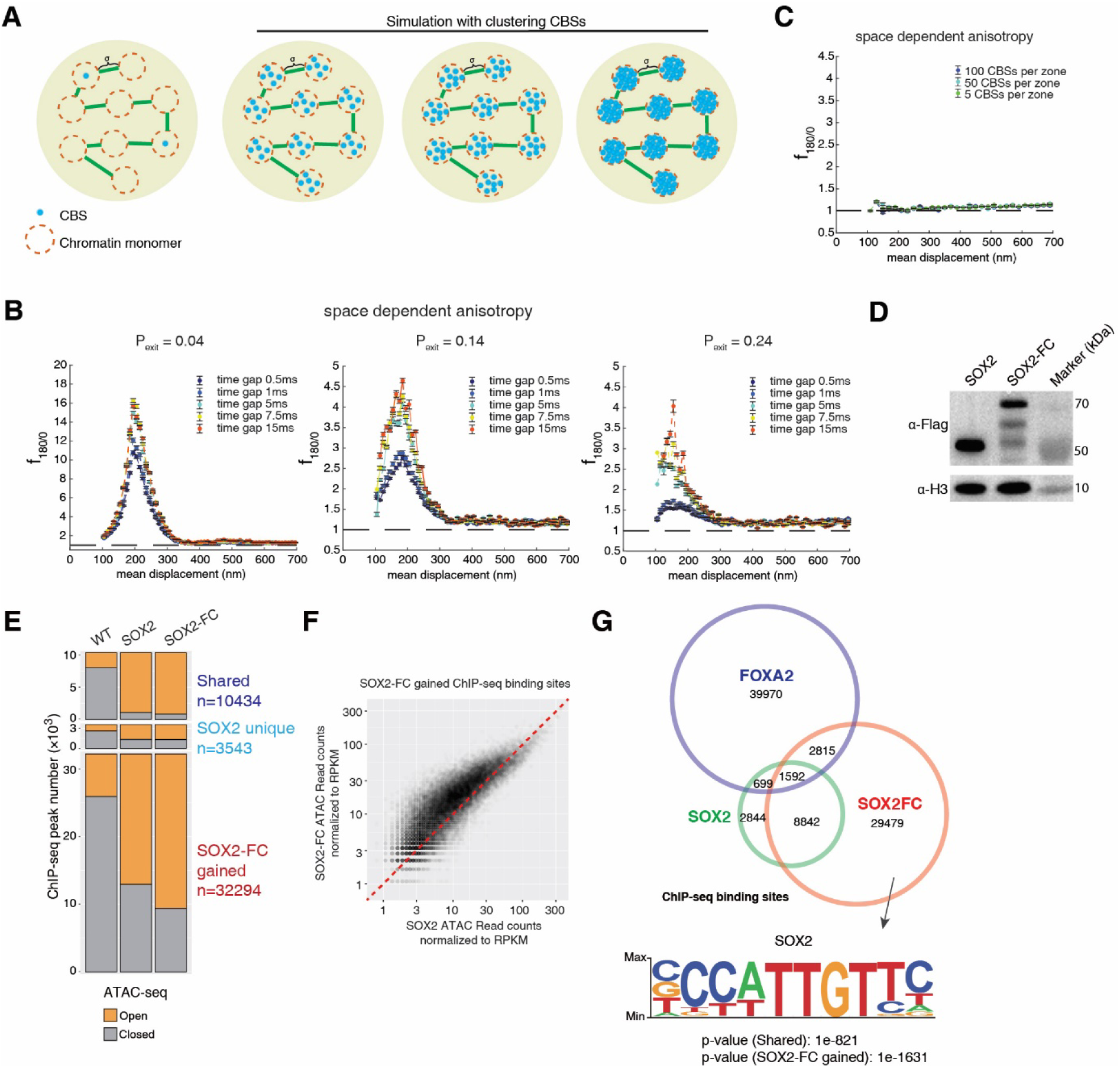
Target search simulation and fusion of CTD_FOXA1/2_ to other TFs. (**A**) Chromatin is simulated as a chain of N monomers connected by harmonic springs. In the first scenario, a small fraction of monomers is randomly chosen to contain CBS (orange dashed-circles with a blue circle representing CBS in the middle) and the rest do not. The distance between monomers reaches final equilibrium via Lennard-Jones (LJ) forces with the characteristic LJ distance σ. In the second scenario, chromatin is simulated as before excepting that each monomer is filled with a defined number of CBS. (**B**) Space-dependent anisotropy plots using the simulated trajectories at indicated time gap and P_exit_. (**C**) Space-dependent anisotropy plots using the simulated trajectories at the indicated number of CBS per monomer. (**D**) Western blots of the whole cell lysates from the indicated cell lines with indicated antibodies (repeated twice independently with similar results). (**E**) Numbers of ChIP-seq peaks in either open or closed chromatin based on ATAC-seq. (**F**) Read counts comparison of ATAC-seq signal at SOXA2-FC gained ChIP-seq peaks. (**G**) Venn plots of ChIP-seq peaks (top), and p-values of SOX2 motif enrichment from HOMER analysis of binding sites in the group of “shared” and “SOXA2-FC gained” (down). Significance was calculated using one-way ANOVA followed by Tukey’s multi-comparison, *p-value<0.05, **p-value<0.01, ***p-value<0.001, n.s., not significant.

**Supplementary Table 1.**
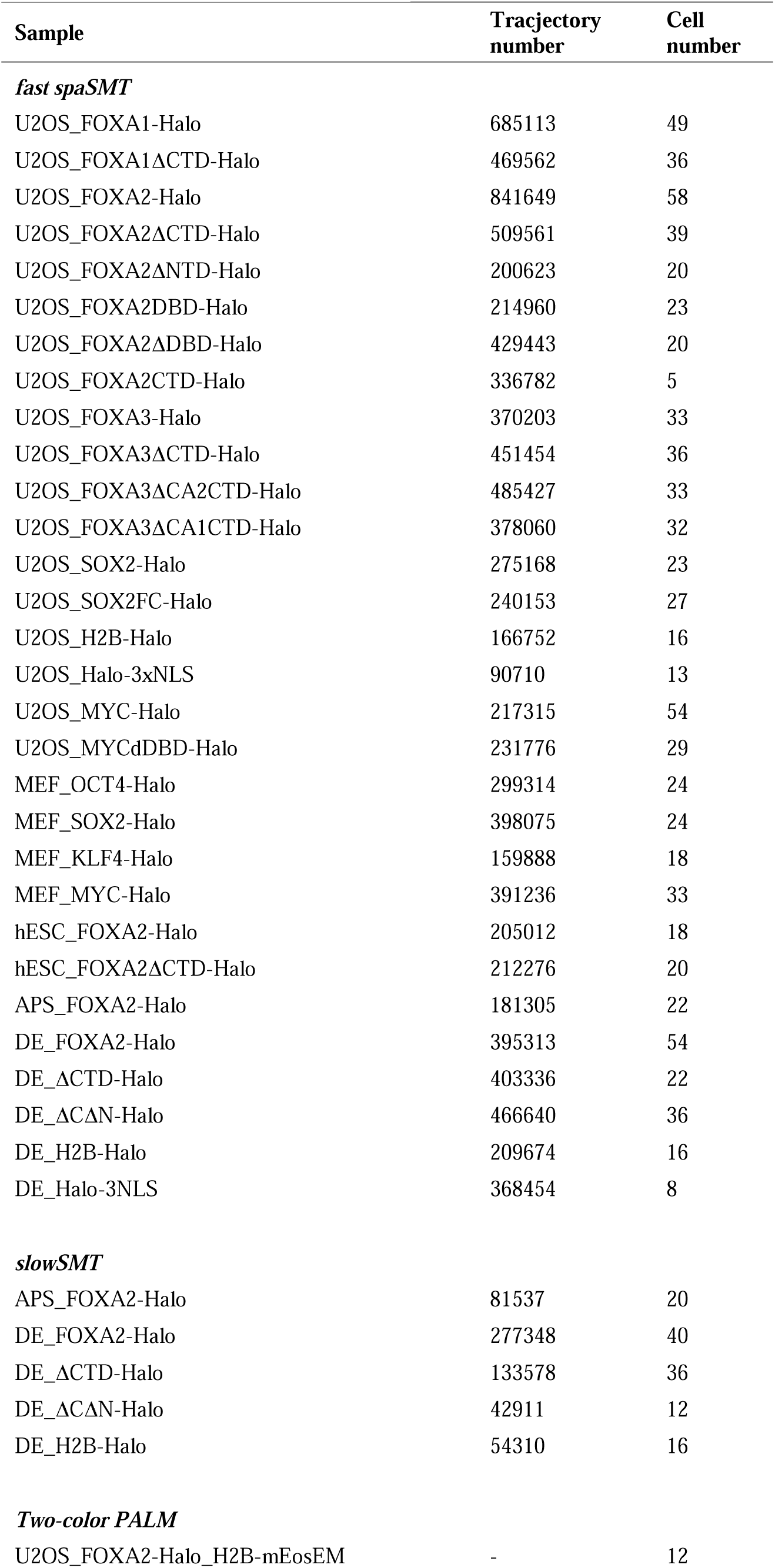
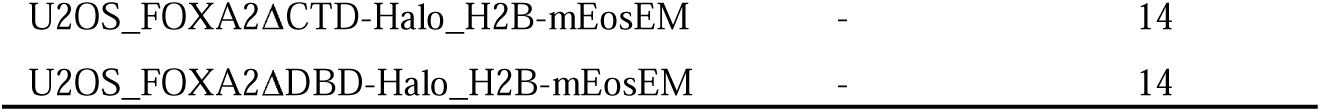

**Supplementary Table 2.**

|  | vbSPT (hidden markov model) |  |  |  | Spot-On |  |
| --- | --- | --- | --- | --- | --- | --- |
| | Slow component | | Fast component | | | Diffusion coefficient of free molecules (mean±s.d., $\mu\text{m}^2/\text{s}$) |
| SampleName | Diffusion coefficient ($\mu\text{m}^2/\text{s}$) | Fraction | Diffusion coefficient ($\mu\text{m}^2/\text{s}$) | Fraction | Bound fraction (mean±s.d.) | |
| U2OS_H2B-Halo | 0.1807 | 0.888 | 4.893 | 0.112 | 0.814±0.079 | 4.123±1.562 |
| U2OS_FOXA1-Halo | 0.1766 | 0.795 | 4.133 | 0.205 | 0.591±0.071 | 2.277±0.534 |
| U2OS_FOXA2-Halo | 0.1937 | 0.808 | 6.094 | 0.192 | 0.551±0.076 | 3.28±0.435 |
| U2OS_FOXA2ΔDBD-Halo | 0.2714 | 0.272 | 7.371 | 0.728 | 0.097±0.035 | 5.875±0.551 |
| U2OS_MYC-Halo | 0.2849 | 0.365 | 5.871 | 0.635 | 0.187±0.043 | 4±0.661 |
| U2OS_MYCΔDBD-Halo | 0.4645 | 0.239 | 7.216 | 0.761 | 0.1±0.036 | 5.259±0.716 |
| U2OS_Halo-3xNLS | 0.2369 | 0.221 | 6.720 | 0.779 | 0.079±0.016 | 5.922±0.839 |
| MEF_OCT4-Halo | 0.1989 | 0.561 | 5.952 | 0.439 | 0.286±0.072 | 4.291±0.98 |
| MEF_SOX2-Halo | 0.2047 | 0.727 | 4.280 | 0.273 | 0.437±0.068 | 2.325±0.58 |
| MEF_KLF4-Halo | 0.2261 | 0.452 | 5.616 | 0.548 | 0.216±0.053 | 4.551±0.832 |
| MEF_MYC-Halo | 0.3616 | 0.320 | 6.043 | 0.680 | 0.125±0.049 | 4.172±1.115 |

**Supplementary Table 3.**

| Cell type | HaloTag fusion | Total bound fraction (%) | Fraction of specific binding in all binding events<sup>#</sup> (%) | $N_{trial}$ (the number of trials to find a specific site) | Specific residence time, RT<sub>s</sub> (s) | Nonspecific residence time, RT<sub>ns</sub> (s) | $\tau_{free}$ (s) | $\tau_{search}$ (s) | Observed association rate on specific targets, $k_{on,s}^*$ (s<sup>-1</sup>) | Time partition of free diffusion in target search (%)<sup>##</sup> |
| --- | --- | --- | --- | --- | --- | --- | --- | --- | --- | --- |
| DE | FOXA2-Halo | 58.1±1.4 | 10.0±0.2 | 167±21 | 103.2±11.9 | 1.76±0.03 | 1.65±0.03 | 557.0±76.6 | 0.0018±0.0002 | 49.50% |
| DE | ΔCTD-Halo | 37.6±2.3 | 9.1±0.2 | 188±57 | 151.1±33.5 | 1.72±0.03 | 4.19±0.06 | 1113.3±337.6 | 0.0010±0.0002 | 71.80% |

Note:
Data are presented as mean ± S.D.
#: the fraction of specific binding is calculated by slowSMT data (see material and methods)

**Supplementary Table 4.**
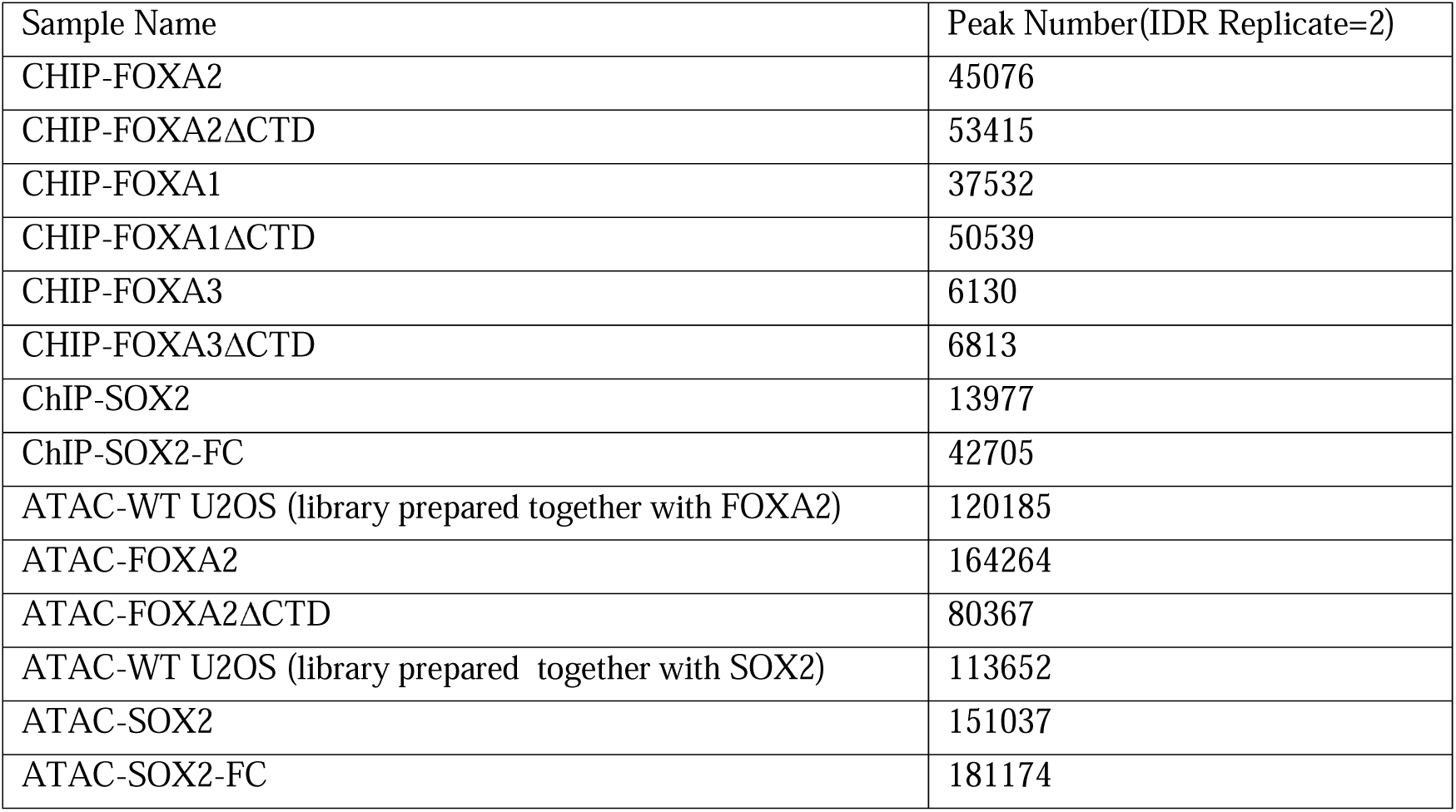

| Sample Name | Peak Number(IDR Replicate=2) |
| --- | --- |
| CHIP-FOXA2 | 45076 |
| CHIP-FOXA2ΔCTD | 53415 |
| CHIP-FOXA1 | 37532 |
| CHIP-FOXA1ΔCTD | 50539 |
| CHIP-FOXA3 | 6130 |
| CHIP-FOXA3ΔCTD | 6813 |
| ChIP-SOX2 | 13977 |
| ChIP-SOX2-FC | 42705 |
| ATAC-WT U2OS (library prepared together with FOXA2) | 120185 |
| ATAC-FOXA2 | 164264 |
| ATAC-FOXA2ΔCTD | 80367 |
| ATAC-WT U2OS (library prepared together with SOX2) | 113652 |
| ATAC-SOX2 | 151037 |
| ATAC-SOX2-FC | 181174 |

**Supplementary Table 5.**
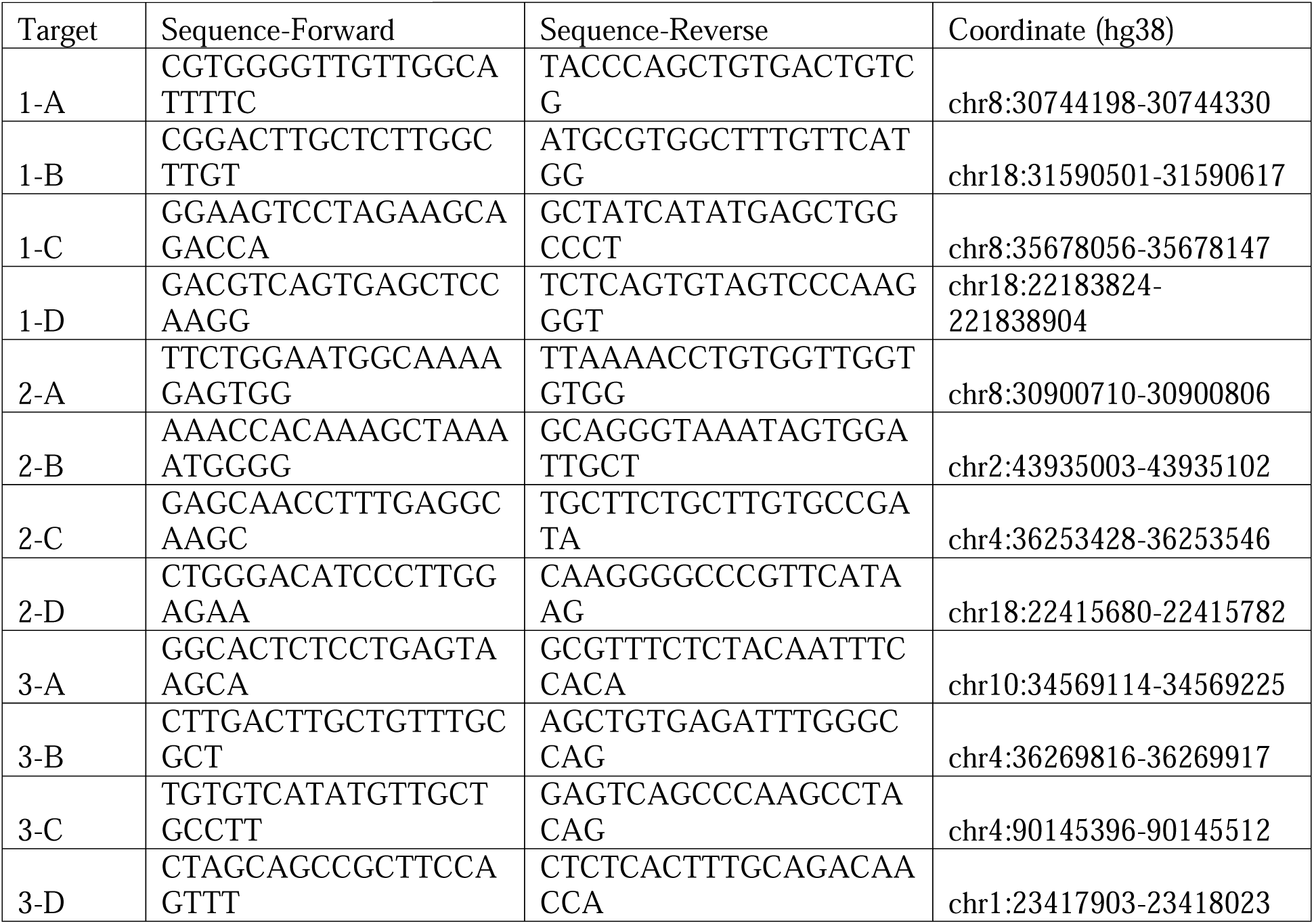

**Supplementary Table 6.** The document is too large to fit here, please see the attached file. Note: p-value was calculated by the Wald test implemented by DESeq2 with Benjamini-Hochberg adjustment.

**Supplementary Table 7.**

| Parameters | Description | Value |
| --- | --- | --- |
| R | Radius of the simulated nucleus | 5 $\mu\text{m}$ |
| $\Delta t$ | Minimal time gap of the simulation | 0.1 msec |
| $l_0$ | Polymer persistence length | 600 nm |
| $\sigma$ | Monomer-monomer Lennard-Jones (LJ) potential equilibrium distance | Same as $l_0$, 600 nm |
| $k$ | Spring constant | $\frac{3k_B T}{(0.2l_0)^2} \text{ Nm}^{-1}$ |
| N | Total number of the monomers | 450 |
| $f_{\text{CBS}}$ | Fraction of the monomers with a cognate binding site (CBS) | 0.02 |
| $D_1$ | Diffusion constant of the free protein when untrapped | 5 $\mu\text{m}^2/\text{sec}$ |
| $D_2$ | Diffusion constant of the free protein when trapped | 2.5 $\mu\text{m}^2/\text{sec}$ |
| $\epsilon_{\text{mono}} / \epsilon_{\text{CBS}}$ | Trapping radius of the monomer / the CBS | 200 nm / 30 nm |
| $\alpha_{\text{mono}} / \alpha_{\text{CBS}}$ | Releasing radius of the monomer and CBS | 210 nm / 40 nm |
| $P_{\text{enter}}$ | Trapping probability to the monomer when hitting the boundary from the outside of the monomer | 0.99 |
| $P_{\text{exit}}$ | Exit probability from the monomer when hitting the boundary from the inside of the monomer | 0.14~0.54 |
| $k_{\text{on}}$ | On-rate to bind nonspecifically inside a monomer | 70 $\text{sec}^{-1}$ |
| $k_{\text{off}}$ | Off-rate to be released from a nonspecific binding site inside a monomer | 200 $\text{msec}^{-1}$ |
| $\tau_{\text{CBS}}$ | Specific binding time in CBS. Randomized from a Poisson distribution with mean $\tau_{\text{CBS}}$ | 12.5 sec |

**Supplementary Table 8.**

| Name | Mean of<br>f180/0,<br>200nm | P_exit | Fraction of<br>time in<br>confined<br>diffusion | Consecutive<br>binding on CBS<br>in the same<br>monomer | Apparent k_on<br>(s<sup>-1</sup>) |
| --- | --- | --- | --- | --- | --- |
| U2OS FOXA1-Halo | 2.94 | 0.209 | 0.2 | 2.89 | 0.0808 |
| U2OS FOXA2-Halo | 3.34 | 0.174 | 0.227 | 3.18 | 0.0929 |
| MEF OCT4-Halo | 2.44 | 0.272 | 0.168 | 2.56 | 0.0663 |
| MEF SOX2-Halo | 3.17 | 0.187 | 0.216 | 3.06 | 0.0879 |
| MEF KLF4-Halo | 1.7 | 0.453 | 0.118 | 2.13 | 0.0449 |
| MEF MYC-Halo | 1.17 | 0.77 | 0.082 | 1.87 | 0.03 |
| U2OS MYC-Halo | 1.19 | 0.756 | 0.083 | 1.88 | 0.0304 |
| U2OS Halo-3xNLS | 1.21 | 0.736 | 0.085 | 1.89 | 0.0311 |
| U2OS FOXA2ΔCTD-Halo | 2.27 | 0.301 | 0.156 | 2.45 | 0.0613 |
| U2OS FOXA2ΔDBD-Halo | 1.4 | 0.598 | 0.098 | 1.98 | 0.0364 |

**Supplementary Table 9.**
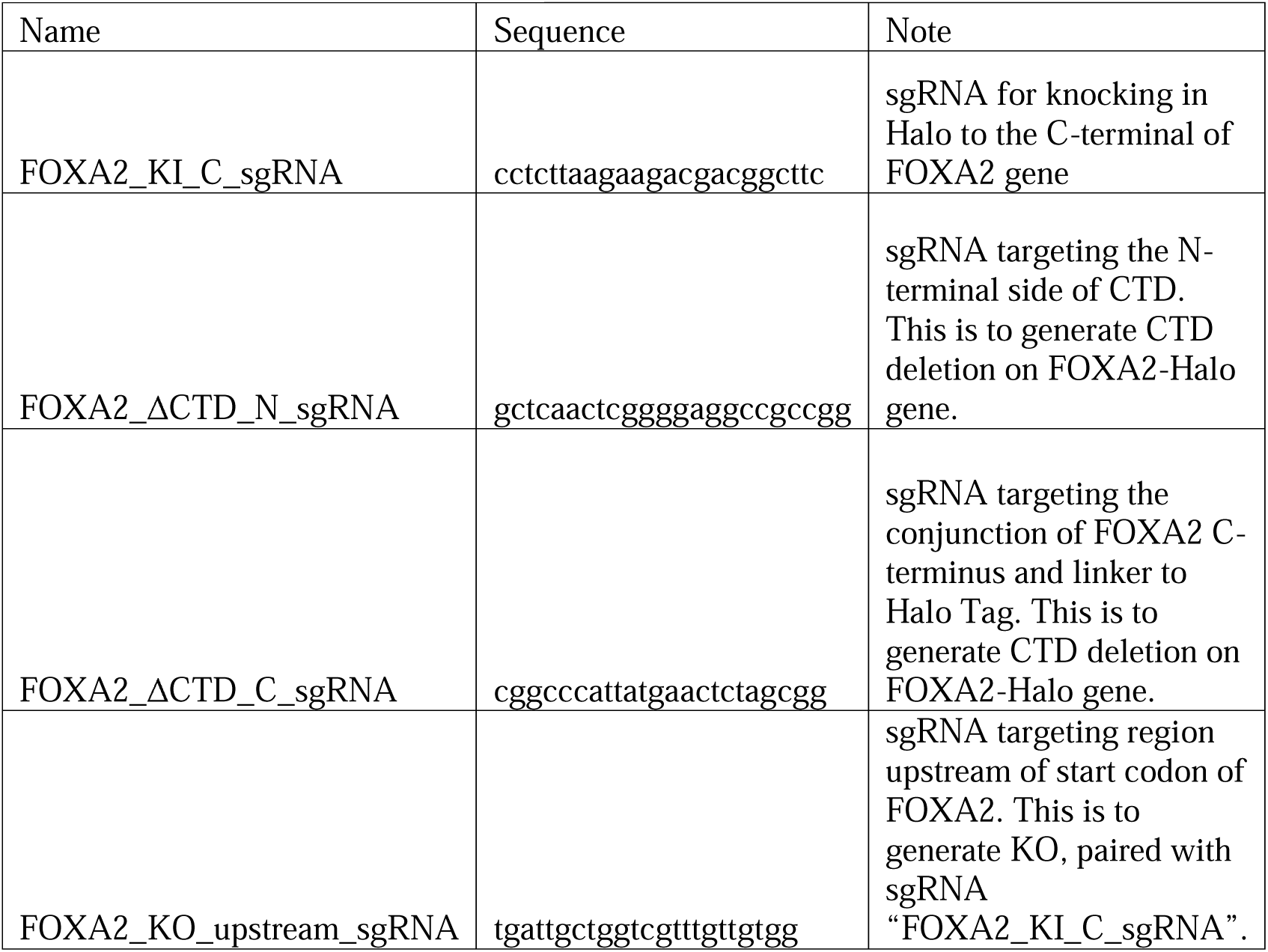

**Supplementary Table 10.**

| Name | Catalog # | Usage | Dilution |
| --- | --- | --- | --- |
| anti-FOXA2 | R&D, AF2400-SP | Western blot | 1:500 |
| anti-SOX17 | R&D, AF1924-SP | Western blot | 1:1000 |
| anti-Halo | Promega, G9211 | Western blot | 1:1000 |
| anti-TBP | Abcam, ab51841 | Western blot | 1:1000 |
| anti-H3 | Beyotime, AH433 | Western blot | 1:2000 |
| anti-H4 | Beyotime, AH458 | Western blot | 1:500 |
| anti-Flag | Sigma, A01868-40 | Western blot and Co-IP | 1:1000 |
| anti-Flag | Sigma, F1804 | ChIP-seq | 1:1000 |
| Goat Anti-Rabbit IgG (H&L)-HRP Conjugated secondary antibody | EASYBIO, BE0101 | Western blot | 1:10000 |
| Goat Anti-Mouse IgG (H&L)-HRP Conjugated secondary antibody | EASYBIO, BE0102 | Western blot | 1:10000 |
| Rabbit Anti-Goat IgG (H&L)-HRP Conjugated secondary antibody | EASYBIO, BE0103 | Western blot | 1:10000 |

## Methods

### Cell culture and stable cell line

U2OS cells, HEK293T cells and MEF cells were cultured in the complete DMEM medium (Dulbecco’s Modified Eagle Medium (Gibco, C11995500BT) supplemented with 10% fetal bovine serum (VisTech, SE100-011) and 1% penicillin/streptomycin (Hyclone, SV30010)). The stable cell lines of U2OS cells with the Tet-on inducible system were constructed with the PiggyBac Transposon system (System Biosciences) containing a puromycin selection gene. Briefly, the Tet-on plasmid and supertransposase plasmid were co-transfected into U2OS cells using lipofectamine 3000 (Thermo Fisher, L3000075) according to the manufacturer’s instructions. 48 hours post transfection, 1 ug/ml puromycin (Selleck, S7417) were added to the medium to select for positively integrated cells.

Lentivirus Tet-on inducible lentiviral system was packaged in HEK293T cells with standard protocol of Calcium phosphate transfection, collected at 36 hours post-transfection and filtered through 0.45-μm pore size filters (Pall, 4614). To image infected MEF cells, 2×10^4^ early-passage MEF cells were first seeded on a 4-well Cellvis chamber (Cellvis # C4-1.5H-N) for 24 hours before viral infection, then infected with viral supernatant of OCT4-Halo, SOX2-Halo, KLF4-Halo, or MYC-Halo, rtTA viral supernatants and 8 μg/ml polybrene (MacGene, MC032). The viral-containing medium was replaced with the fresh medium 12 hours post-infection and Dox were added at 24 hours post-infection. MEF cells were stained and imaged 48 hours post infection. All plasmids are available upon request.

### hESC culture, differentiation and genome editing

H7 hESCs (WiCell, WAe007-A) were cultured on plates pre-coated with 1% (v/v) Geltrex (Thermo Fisher, A1413302) with complete Essential 8 medium (Thermo Fisher, A1517001). Accutase (Thermo Fisher, A1110501) was used for cell dispersion and ROCK inhibitor Thiazovivin (Tocris, 3845) at a final concentration of 1 μΜ was added to the culture medium for the first day after re-plating cells. hESCs were differentiated to APS and DE with a two-day protocol using PSC Definitive Endoderm Induction Kit (Thermo Fisher, A3062601) or home-made media ^36^. Cells were stained with surface marker APC-CXCR4 (BD 555976) at a 1:5 dilution for 30 minutes on ice and examined by flow cytometry to assess differentiation rate.

For genome editing, sgRNAs were designed using CRISPOR (http://crispor.tefor.net/crispor.py) and cloned into a plasmid expressing Cas9 and Venus fluorescent protein. Sequences for genome editing sgRNAs are listed in Supplementary Table 9. hESCs were transfected using Fugene 6 (Promega, E2691) following manufacturer instructions. For knock-in, a Cas9/sgRNA plasmid and a donor plasmid were co-transfected; for knock-out or domain deletion, two Cas9/sgRNA plasmids were co-transfected. 48 hours post-transfection, Venus positive cells were sorted by FACS, plated at low density, and cultured for about 10 days to generate discrete cell clones. Individual cell clones were picked out and genotyped by PCR to identify desired homozygous clones. The desired clones were further verified by Sanger sequencing and Western blots.

### Microscopy setup

All SMT and PALM experiments were performed on a custom-built Nikon Ti2 microscope as before ^55^. It is equipped with two EM-CCD cameras (Andor, iXon Ultra 897), a 100x/NA 1.49 oil-immersion TIRF objective (Nikon apochromat CFI Apo TIRF 100x oil), a perfect focusing system (Applied Scientific Instrumentation), and a stable-top chamber for maintaining 37℃ and 5% CO2 (Tokai Hit). The HILO illumination was achieved by a Nikon TIRF module, multiple lasers (405 nm: maximum 140mW, OBIS, Coherent; 560 nm: maximum 1 W, MPB; 642 nm: maximum 1.5 W, MPB) controlled by the AOTF system (AA Opto-Electronic, AOTFnC-VIS-TN), a multi-band dichroic mirror (405/488/561/633 nm quad-band, Semrock) and emission filters (TMR/PA-JF549: Semrock 593/40 nm band-pass filter; JF646/PA-JF646: Semrock 676/37 nm bandpass filter).

### Cell labeling for SMT, PALM and confocal imaging

U2OS and MEF cells were plated on Mat-Tek dishes (Mat-Tek # P35G-1.5-14-C) or 8-well Cellvis chambers (Cellvis # C8-1.5H-N) with #1.5 high performance cover glass, and induced protein expression with 1μg/mL doxycycline (Selleck, S4163) for one or four days. For hESCs and their differentiation, the culture dish with glass bottom was first coated with Geltrex (Thermo Fisher, no. A1413301) overnight. To label Halo fusion protein in living cells, Halo ligand conjugated fluorescent dyes at indicated final concentration were added to culture medium (fast spaSMT: 25 nM PA-JF646 or 2.5 nM PA-JF549; slowSMT: 25 pM JF646; PALM: 25 nM PA-JF646; confocal imaging: 200nM TMR) and incubate at cell incubator for 30 min. The medium was then removed and two rounds of incubation with fresh medium for 30 minutes were performed to remove free dyes. Right before live-cell imaging of SMT, the medium was changed to phenol red-free medium. For two-color PALM and confocal imaging of fixed cells, after Halo staining, cells were cross-linked with pre-warmed 4% PFA for 10 min at 37°C, permeabilized with 0.1% v/v Triton X-100 in PBS for 10 min at room temperature, and then washed with PBS. Medium used in each step was pre-warmed and caution was taken to avoid cells to detach from the glass bottom.

### Acquisition of SMT data

The 133 Hz fast spaSMT experimental settings were as follows: in one frame of camera exposure time (7 ms), 1 ms of 633 nm excitation (100% AOTF) was delivered at the beginning of the frame; 405 nm photo-activation pulses were delivered during the camera integration time (∼447 μs) to minimize background and their intensity was optimized to achieve a mean density of 1-2 molecules per frame per nucleus. In addition to 7 ms (∼133 Hz), the camera exposure times were also set at 13 ms (∼74 Hz) to obtain additional data for analyzing the space dependency of anisotropy. 20,000 frames were recorded per cell per experiment. For each sample, we performed two or three independent replicates and recorded data of multiple cells in each replicate.

The slowSMT experiments used a long exposure time (500 ms) and continuous illumination with low laser intensity to capture the position of slow mobile molecules. Under this mode, fast-moving molecules will be blurred into the background noise. Caution was taken to ensure cells did not have gross movement during the process of imaging to minimize tracking error. Generally, each cell was captured for 1500 frames with 500 ms exposure time. For each sample, we performed two or three independent replicates and recorded data of multiple cells in each replicate.

### Acquisition of simultaneous two-color PALM data

A 405 nm laser was used during the camera integration time (∼447 μs) to photoconvert mEosEM and photoactivate PA-JF646 at sparse density. The 561 nm and 642 nm lasers were used to illuminate cells for continuous 30 ms and two colors were detected simultaneously with two EM-CCD at an exposure time of 30 ms. 30,000 frames of images were taken for each cell, and at least twelve cells were recorded for each sample. In combined live-cell PALM and fast spaSMT experiments, PALM data was collected with an exposure time of 30 ms, fast spaSMT data was collected immediately with an exposure time of 7.5 ms.

### Single-molecule localization and trajectory linking

Both fast spaSMT movies and slowSMT movies were analyzed with custom-made MATLAB scripts (https://gitlab.com/tjian-darzacq-lab/SPT_LocAndTrack) implementing a multiple-target tracking (MTT) algorithm ^56^ to generate single-molecule trajectories. The settings used in the MATLAB scripts for fast spaSMT were as follows: localization error, 10^-6.25^; deflation loops, 0; Blinking (frames), 0; maximum competitors, 3; maximum D (μm^2^ s^-1^), 20. The settings used in the MATLAB scripts for slowSMT were as follows: localization error, 10^-6.25^; deflation loops, 0; Blinking (frames), 1; maximum competitors, 3; maximum D (μm^2^ s^-1^), 0.7.

### Estimation of bound fraction and diffusion coefficient from fast spaSMT data

Fraction of bound molecules and diffusion coefficient of unbound molecules were determined by fitting the two-state Spot-On kinetic model ^31^ (v1.05; GitLab tag 6a3e885c) with the spaSMT trajectories obtained with 7.5 ms exposure time. The parameters used in the fitting were as follows: dZ = 0.7μm; GapsAllowed = 1; Time Points = 8; JumpsToConsider = 4; Model Fit = 2; NumberOfStates = 2; FitLocError = 0; LocError = 0.040 μm; D_Free_2State = [0.5, 25]; D_Bound_2State = [0.0001, 0.04].

### Residence time inference and survival analysis of slowSMT data

The trajectory lengths of stably bound molecules from slowSMT experiments were plotted as a survival curve (1-CDF). We adopted the method from ^55^ to calculate residence time of transcription factors on targets.

### Calculation of target search kinetics

To investigate how FOXA2 explores the nucleus, we adopted the method described by ^18^ to derive various target searching parameters.

### Angular analysis of fast spaSMT data

We used the Matlab scripts written from (https://gitlab.com/anders.sejr.hansen/anisotropy) ^38^ with some changes to analyze the distribution of the molecule trajectories obtained from our fast spaSMT experiments. Briefly, we merged trajectories from different cells obtained from the fast spaSMT datasets (133 Hz, and 74 Hz). In the merging step, we also did a quality control step to further ensure that the downstream analysis is not subject to the wrong connection of two different molecules observed in consecutive frames during reconnection steps in the MTT algorithm. “ClosestDist” was set to 2 so that if any two detected localizations in the same frame are within 2 μm their trajectories will be discarded from downstream analysis. The merged trajectories were HMM-classified to label each segment of a trajectory as either “bound” or “free”, and the bound segments of each trajectory were removed. The angles formed by two consecutive “free” segments with minimum length of 100 nm (“GlobalMinJumpThres” was set to 0.1, which stands for 100 nm) were calculated. The angles from all free segments were pooled and the fold of the angles ranging from 180 ± 30° versus 0 ± 30°, f180/0 (i.e., global anisotropy), were calculated to measure the anisotropic behavior of the molecules.

In addition to analyzing angles from all free segments, we also analyzed the angles composed of free segments right before or after one bound segment (i.e., local anisotropy). Specifically, after classifying the segments of each trajectory into “bound” or “free” states by the HMM model as above, we selected a defined number of the consecutive free segments either before or after a bound segment. Then we discarded any “free” segments less than 100 nm to calculate angles between the two adjacent free segments and pooled all angles to calculate the f180/0, local.

### Processing and rendering PALM data

Raw imaging data with single-molecule signals were analyzed using the ImageJ plug-in ThunderStorm ^57^ to detect single molecule localizations. Low-quality single molecule signals (PSF too small or too large or with high uncertainty) were filtered out. Single molecule localization data consisting of x and y localization lists were used to render PALM images, by assigning pixel size as 10 nm and pixel value equal to the number of localizations within the pixel area.

### Two-color PALM data processing and model-based chromatin segmentation

Two-color PALM raw data were analyzed by ThunderStorm to generate molecule localizations and merge reappearing localizations in subsequent frames. The resulted localization data were further analyzed by a self-written MATLAB script. Sample drift was determined by calculating the spatial cross correlation function between sequential subsets of localizations and corrected. Data with major sample drift was discarded to ensure high data quality. We generated rendered super-resolution images with pixel size at 10 nm and pixel value as the number of molecular localizations within the pixel. Manders’ correlation coefficents were calculated by ImageJ Colocalization Threshold plugin using H2B PALM and FOXA2 PALM images as input and default threshold.

A Voronoi diagram was created by the SR-Tesseler analysis ^42^, after which clusters were identified by setting the density factor to 2.5. Clusters were filtered by Min area and Min # locs (Min area:20; Min # locs: 50). The following information was directly exported: area, # detections, circularity, and diameter.

Chromatin compaction levels were determined by the intensity of rendered H2B super-resolution images with 10nm pixel size. The nuclear border was determined by Gaussian smoothing and threshold binarization. And the resulted images were further analyzed by a hidden Markov random field model to segment images to seven classes ^44^ based on H2B intensity.

After the chromatin compaction segmentation into seven classes, localization densities of different FOXA2 variant in each chromatin class were calculated and divided by densities of localizations following a random distribution; the relative density enrichments were presented as heatmaps. The seven chromatin intensity classes were further divided into three biologically relevant regions: interchromatin (IC, class 1), perichromatin (PC, classes 2-3), and chromatin domain (CD, classes 4-7), as described in ^44^. The CD region was composed of chains of individual CDs. Their centroids were found by the regional maximum algorithm and the borders of individual CD were determined by the intensity valley calculated from intensity gradients. The centroids of FOXA2 variants’ clusters were also determined by the regional maximum algorithm, and only centroids that located within an individual CD were selected out and their distance to the corresponding CD centroid were measured.

As a parallel method, the cell nucleus was also divided into nine equal-area parts from low to high density based on the intensity of H2B and FOXA2 variants respectively, and the localization number and percentage of H2B and FOXA2 variants within each H2B region, as well as the overlapped areas between H2B and FOXA2 at different densities were calculated. Heatmaps were plotted with x and y axes representing nine density regions index of FOXA2 variants and H2B from low to high, respectively.

In combined live-cell PALM and fast spaSMT experiments, 1000 frames of images with 30 ms exposure time were used to render a PALM image, and fast spaSMT data were collected immediately after PALM and overlaid.

### Simulation of chromatin and the TF target search

We used a previously developed chromatin model ^38,58^ that treats chromatin as a coarse-grained, self-avoiding polymer confined within the nucleus. Briefly, we modeled the chromatin as a series of *N* monomers with a radius of 200 nm, with positions (*R_1_,…,R_n_*). A small portion of randomly chosen monomers had cognate binding sites (CBSs) placed in their middle (for confined target search model) or filled with the defined number of CBSs randomly (for clustering target sites model). The chromatin chain first reached equilibrium with the defined spring potential and Lennard-Jones forces, then each monomer position was fixed (Supplementary Fig. 6A). All monomers were confined within a sphere with radius *R* to simulate cell nucleus, and each rigid monomer was reflected in the normal direction of the tangent plane of the sphere boundary.

For the simulation of confined target search model, simulated TF interacts with the modeled chromatin in three ways. First, TF underwent free Brownian motion when not interacting with monomers. Second, when TF touched the boundary of monomer (the distance between TF and the center of monomer smaller than ε_mono_), it has a certain probability (P_enter_) of entering the monomer. Once entering the monomer, TF diffused inside the monomer and exited it with certain probability (P_exit_) once hitting the boundary of the monomer. When diffusing inside the monomer, TF can transiently bind with a Poissonian ON-rate *k_on_* and then be released with a Poissonian off-rate *k_off_* to mimic non-specific binding. If the monomer contains a CBS in its middle, TF binds to CBS when its distance to the center of the CBS (same as the center of monomer) is smaller than ε_CBS_. The residence time of CBS binding followed a Poisson distribution with a mean time τ_CBS_ 12.5 sec. The value of residence time did not affect the simulation results, since we only focused on target search, i.e. diffusion, rather than target binding part of the final trajectory. After finishing binding on CBS, TF continued diffusing inside the monomer. Of note, we used Euler’s scheme to generate the Brownian motion of the protein.

In the simulation of the clustering target sites model, the TF undergoes free Brownian motion when not interacting with CBSs. The TF binds to CBS when its distance to the center of the CBS is smaller than εCBS. The residence time of CBS binding follows a Poisson distribution with a mean time of τCBS, which is 12.5 seconds. After completing the binding on CBS, the TF continues to diffuse inside the nucleus.

Each round of simulation generated a long Brownian trajectory with the time gap of 0.1 ms. The protein was placed at release radius (α_mono_) of a randomly chosen monomer and started diffusing. Each round of simulation stopped when total steps reached 5,000 or when bound to CBS for 500 times, depending on the downstream analysis. To simulate the localization error of real imaging conditions, white Gaussian noise with a standard deviation of 30 nm and zero mean was added to each trajectory point. The generated trajectories were then projected into 2D (discarding Z position) and split into short trajectories with the desired time gap (e.g., 7.5 ms) to mimic the exposure time used in fast spaSMT. We performed 12 simulations for each condition. The generated trajectories were then pooled and analyzed using the same pipeline as in “Angular analysis of fast spaSMT data”. More detailed parameters used in the simulation were listed in Supplementary Table 7.

### Co-immunoprecipitation

Stable HeLa cell lines were induced with 1 μg/ml doxycycline for 48 hours, collected and gently lysed with ice-cold NP-40 lysis buffer (10 mM HEPES pH 7.5, 10 mM KCl, 0.2 mM EDTA, 1 mM DTT, 0.5% NP-40) on ice for 20 min. The pelleted nuclei were further extracted with high salt buffer (500 mM NaCl) by incubation on ice for 60 min. The nuclear lysate was immunoprecipitated with pre-activated Flag-Magnetic Dynabeads (Thermofisher, 88847) at 4°C overnight, followed by one wash with balance lysis buffer (25 mM HEPES pH 7.5, 0.2 mM EDTA, 0.5% NP-40), three washes with high-salt wash buffer (25 mM HEPES pH 7.5, 500 mM NaCl, 0.2 mM EDTA, 0.5% NP-40), one wash with low-salt wash buffer (25 mM HEPES pH 7.5, 150 mM NaCl, 0.2 mM EDTA, 0.5% NP-40), and boiling in SDS-PAGE loading buffer at 100°C for 10 min. The antibodies used were listed in Supplementary Table 10.

### ChIP-seq and ATAC-seq

The ChIP-seq followed the standard protocols ^59^. ChIP-seq experiments were performed in two replicates. Primary antibodies for ChIP-seq were listed in Supplementary Table 10.

OmniATAC-seq was performed according to the protocol from ^60^. ATAC-seq experiment was performed in two replicates.

For ChIP-seq and ATAC-seq, raw reads were quality-checked with FastQC (version 0.10.1) and trimmed by trim_galore function (v0.6.10), with --paired mode and the default parameters. Reads were aligned to human genome (hg38) using bowtie2 (v2.4.4). The resulted sam files were converted to bam files with samtools (v.0.1.18). The bam files of the two replicates of one treatment were called peaks individually using MACS2(v2.2.7.1) with the default parameters, and the resulted narrowPeak files were used to call peaks by the IDR(2.0.4.2) method. The bamCoverage function in deeptools(v2.4.1) was used to convert bam files to bigwig files for visualization, and normalized by RPKM. A high correlation above 0.8 was observed between replicates for all samples. The R package intervene (v0.6.5) was used to determine overlaps among two or more peak sets. The function computeMatrix in the deepTools suite (v.2.4.1) was used with reference-point mode to calculate scores for each genomic region. Motif analysis of differential peaks from each group were performed by software HOMER (v4.11). Genome-wide chromatin state predictions for each biological condition were performed using ChromHMM (v1.24) software with default parameters. To compare the ChIP-seq intensity from FOXA2 and FOXA2 ΔCTD, ChIP-seq libraries were normalized to input-calibrated ChIP-qPCR of 12 sites that comprised varying signal intensities, using a linear regression model (Supplementary Fig. 4B, Supplementary Table 5) as in ^61^.

### RNA-seq

Poly-A RNA-seq was performed for undifferentiated hESC or hESCs differentiated to APS and DE. The purified total RNA was added to ERCC RNA Spike-In Mix 2 (ThermoFisher Scientific # 4456740). Messenger RNA purification, RNA fragmentation, first and second strand cDNA synthesis were performed according to the TruSeq RNA Sample Preparation v2 Kit (Illumina, RS-122-2001) but using Superscript III for reverse transcription instead of Superscript II (50 °C for 50’ incubation time). cDNA was purified with AMPure XP beads (Beckman, A63882) and subjected to a Solexa rapid library protocol. PCR reactions were cleaned up once with AMPure XP beads. Library quality and fragment size were assessed by qPCR and Fragment analyzer™ and sequenced on the Illumina HiSeq4000 sequencing platform (single end-reads, 50 bp long). RNA-seq experiments were performed in two replicates.

### Analysis of RNA-seq data

Reads were quality-checked with FastQC and aligned to the transcriptome of human genome (hg38) by STAR v2.7.1a with following parameters: --outSJfilterReads Unique --outFilterMultimapNmax 1 --outFilterIntronMotifs RemoveNoncanonical --outSAMstrandField intronMotif. The gene expression level was calculated by HTSeq v2.0.2 based on the gencode hg38 annotation (GENCODE v32) with parameters: htseq-count --stranded=no -f bam --additional-attr=gene_name -m union. All subsequent analyses were performed in R.

Differential gene expression analysis was performed on three cell stages (hESC, APS, and DE), with two replicates using DESeq2 v1.30.1. Stage-specific genes were selected with log2(fold change) > 1 and FDR <0.05. FDR was calculated by the Wald test implemented by DESeq2 with Benjamini-Hochberg adjustment. PCA analysis of differentially expressed genes was performed using DESeq2. Putative FOXA2 target genes were identified as genes having ±5kb FOXA2 binding sites flanking the gene body at the DE stage using clusterProfiler v3.18.1. DE stage-specific genes were further sorted based on gene expression changes after FOXA2 knockout and their dependency on CTD deletion using DESeq2-normalized RNA-seq read counts. Genes with the fold change > 1.5 after FOXA2 deletion were classified as “repressed” by FOXA2, while genes with the fold change < 0.66 were classified as “activated” by FOXA2. Within these FOXA2-dependent genes (repressed or activated), their dependency on CTD was further assessed to identify the “CTD-dependent” genes. Therefore, these genes are classified to five groups: I (CTD-dependent gene repression with FoldChange(KO/WT) > 1.5 and FoldChange(ΔCTD/WT) > 1.5), II (CTD-independent gene repression with FoldChange(KO/WT) > 1.5 and 2/3 < FoldChange(ΔCTD/WT) < 1.5), III (CTD-dependent gene activation with FoldChange(KO/WT) < 2/3 and FoldChange(ΔCTD/WT) < 2/3), IV (CTD-independent gene activation with FoldChange(KO/WT) < 2/3 and FoldChange(ΔCTD /WT) > 2/3) and IV (unchanged with 2/3 < FoldChange(KO/WT) < 1.5). Gene Ontology (GO) term enrichment analysis was performed using clusterProfiler and GO terms with a p-value < 0.05 were considered significantly enriched. All volcano plots and bubble charts were plotted using ggplot2, and the heatmap was generated using pheatmap.

## Data Availability

All data are available to reproduce the results shown in the main text or the supplementary materials. Imaging data are available at Zenodo (accession number 10427790). Genomic data have been deposited in NCBI’s Gene Expression Omnibus (accession number: GSE203650).

## Code Availability

The analysis code is deposited at Github (https://github.com/denglabpku/).

## Notes

https://zenodo.org/records/10427790

### Competing Interest Statement

The authors have declared no competing interest.

